# Utilising an in silico model to predict outcomes in senescence-driven acute liver injury

**DOI:** 10.1101/2023.10.11.561528

**Authors:** Candice Ashmore-Harris, Evangelia Antonopoulou, Rhona E. Aird, Tak Yung Man, Simon M. Finney, Annelijn M. Speel, Wei-Yu Lu, Stuart J. Forbes, Victoria L. Gadd, Sarah L. Waters

## Abstract

Currently liver transplantation is the only treatment option for liver disease, but organ availability cannot meet demand and transplant recipients require lifelong immunosuppression. The identification of alternative treatments, e.g. cell therapies, able to tip resolution of injury from inflammation to regeneration requires an understanding of the host response to the degree of injury. We adopt a combined *in vivo-in silico* approach and develop a mathematical model of acute liver disease able to predict the host response to injury. We utilise the Mdm2^fl/fl^ mouse model together with a single Cre induction through intravenous injection of the hepatotropic Adeno-associated Virus Serotype 8 Cre (AAV8.Cre) to model acute liver injury. We derive a complementary ordinary differential equation model to capture the dynamics of the key cell players in the injury response together with the extracellular matrix. We show that the mathematical model is able to predict the host response to moderate injury via qualitative comparison of the model predictions with the experimental data. We then use the model to predict the host response to mild and severe injury, and test these predictions *in vivo*, obtaining good qualitative agreement.

## Introduction

Orthotopic liver transplantation remains the only curative treatment for fulminant and endstage disease, but the availability of donor organs is consistently exceeded by patient requirements, with a mortality rate of over two million deaths worldwide per year^1^. Alternative therapies capable of promoting liver regeneration are urgently needed to support this growing patient burden.

Liver regeneration is a complicated, multifaceted process which is dependent on the nature and burden of injury. Commonly, a multicellular interplay of different spatiotemporally activated pro-inflammatory and pro-regenerative responders contribute to injury resolution and tissue regeneration, with dynamic changes in immune cells, extracellular matrix (ECM) composition and vascularisation of the liver occurring during the repair process. Additionally, the level of senescence in the injured liver is widely considered to influence regenerative success. In fact, hepatocyte senescence, characterised by induction of genes encoding p53 (TRP53), p21 (WAF1) and p16 (INK4A), is a common feature of various human liver diseases including steatosis, acute and chronic injury^2–6^. Corresponding acute and chronic murine liver injury models demonstrate complementary senescence expression profiles to their human counterparts^2,4,7^. Despite this, an understanding of how the degree of senescence impacts key components of the liver injury microenvironment, and how these components change over time remains lacking. Such mechanistic understanding has the potential to improve responses to candidate alternative therapies (e.g. novel cell and drug approaches such as senolytic therapies), ultimately enabling optimisation of treatment outcomes, for example through stratification of patients, identification of intervention timepoints, and determination of preconditioning requirements.

The complexity of the liver injury response makes it challenging to identify, using experimental approaches alone, which microenvironmental changes or cellular responses are key to improving therapeutic interventions, not least because significant numbers of animal studies are required to inform this using conventional analysis methods. The ability to predict which changes will occur, the level of response, and which components are important in mediating injury resolution, relative to the degree of (induced) senescence, would substantially reduce the number of animal experiments required, thereby reducing costs of preclinical studies whilst also providing insights into key injury/regeneration mechanisms.

In this paper we adopt a synergistic *in vivo* and mechanistic mathematical modelling approach to provide mechanistic understanding of the complex biological interactions underpinning regenerative processes. We use the mathematical model to predict the biological response to a given injury challenge that is then tested *in vivo*. Furthermore, *in silico* models can provide predictions for the dynamic behaviour of the variables as a function of continuous time, whereas experimental methods return data at discrete timepoints. Such high fidelity numerical data can be used to provide further insights into the system behaviour, such as the time taken for the induced senescence to be cleared from the system and can be exploited to optimise therapeutic outcomes. For a review of how mechanistic mathematical models can be exploited to overcome translational bottlenecks in driving cell therapies from bench to beside, as well as in the field of regenerative medicine more widely, see Ashmore-Harris et al[NPJREGENMED-01618 (under review)] and Waters, Schumacher and El Haj^8^.

Initial mechanistic mathematical model development requires interrogation of experimental observations of the biological system to allow hypotheses for the causal mechanisms underpinning the system behaviour to be identified, as well as experimental data for model calibration and validation. The depth of *in vivo* data presented here presents an exciting opportunity to build a predictive mathematical model.

Previous studies used the AhCre murine double minute 2 (Ah^Cre^Mdm2^flox/flox^, herein *Mdm2*) mouse strain to induce acute, p21-dependent senescent injury in hepatocytes via administration of the xenobiotic chemical β-naphthoflavone (βNF). This mouse combines expression of the rat Cyp1A1 promoter upstream of Cre recombinase with loxP flanked transgenic *Mdm2*, allowing selective deletion of *Mdm2* in hepatocytes following Cre recombinase expression^2,7^. Mdm2 is a key negative regulator of p53, which positively regulates p21 expression.

In this study we instead administer single dose concentrations of AAV8-TBG-Cre allowing liver specific uptake, and tighter control of the level of Cre recombinase expression in hepatocytes, therefore fine tuning the level of induced senescence. We perform a detailed quantitative histological characterisation of the key players in the regenerative response and report this relative to the level of senescence induced. Motivated by cell-to-cell or cell-to-matrix interactions in the pathogenesis of liver fibrosis we specifically consider the macrophage response, myofibroblast activation, changes in the extracellular matrix (ECM, collagen-I production) and the endothelial/angiogenesis response.

This depth of detailed quantitative experimental data presents a timely opportunity to develop a predictive theoretical model. Continuum mathematical models have been successfully developed for a number of regenerative medicine scenarios^9^. Particularly pertinent are wound healing and liver fibrosis studies, such as those explored previously by Friedman and Hao, who generated partial differentiation equation models for important inflammatory and regenerative responses in liver fibrosis, and described changes in key cell types and cytokines within an assumed region of the liver^10^. To date, no models have been developed which describe the changing nature of the liver microenvironment as a result of or in relation to acute, senescence driven liver injury despite the widespread prevalence of senescence in human liver diseases.

In this paper we detail the ordinary differential equation (ODE) mathematical model that captures acute senescence mediated injury development and is capable of identifying the driving mechanisms leading to inflammation, and show that it qualitatively captures the *in vivo* dataset corresponding to a moderate injury level. We then use the theoretical model to qualitatively predict the response of the injured tissue niche to more severe and milder injury levels and demonstrate excellent qualitative agreement with the *in vivo* data, providing validation of our mechanistic mathematical model.

## Results

### Administration of AAV8.TBG.Cre in Mdm2 mice results in transient upregulation of p21 expression

To determine how senescence influences the regenerative response over time we initially induced senescence in *Mdm2* mice through administration of a single ‘moderate’ dose of 4.16x10^10^GCU of hepatotropic AAV8.TBG.Cre (*c*.*f*. materials and methods **Fig. 1A**). Our data administering this dose to Rosa26^LSL-TdTomato^ mice suggests this induces recombination in **93%** of hepatocytes, based on quantification of the proportion of TdTomato^+^ hepatocytes in the parenchyma (**Supplementary Fig.1, group 3**). The proportion of senescent hepatocytes in the *Mdm2* strain is dynamic, with paracrine signalling from recombinant senescent cells inducing senescence in a proportion of neighbouring cells in tandem with the death and clearance of highly senescent cells^3,8^. For simplicity, we do not distinguish between senescence due to recombination or paracrine signalling. To examine the total senescence level we analysed mRNA expression of the senescence marker p21 from total liver extracts by qPCR (**Fig. 1B**) and confirmed expression was hepatic by histological staining (**Fig. 1C**). We previously found that recombination in *Mdm2* mice, through the AhCre system as a result of βNF, induces p21 expression within two days of administration, with a corresponding statistically significant elevation of serum markers for liver injury within six days^3,8^. Here, we analysed senescence marker expression in liver tissue of mice three, seven and fourteen days after AAV8.TBG.Cre administration (D3, D7, D14, **Fig. 1A**). Results showed a significant increase in p21 expression at D3 and D7 (peak expression at D7, mean fold change 114.6±31.53), with expression comparable to healthy age matched control mice by D14 post induction (**Fig. 1B**). These results are supported by comparable temporal trends in p53 expression (**Supplementary Fig.2A**), indicating that “peak injury” is established by D7 with regeneration/resolution by D14. In contrast, there was no upregulation of p16 expression (**Supplementary Fig.2B**) confirming that senescence in this model is driven by the p53/p21 axis.

**Fig. 1.**
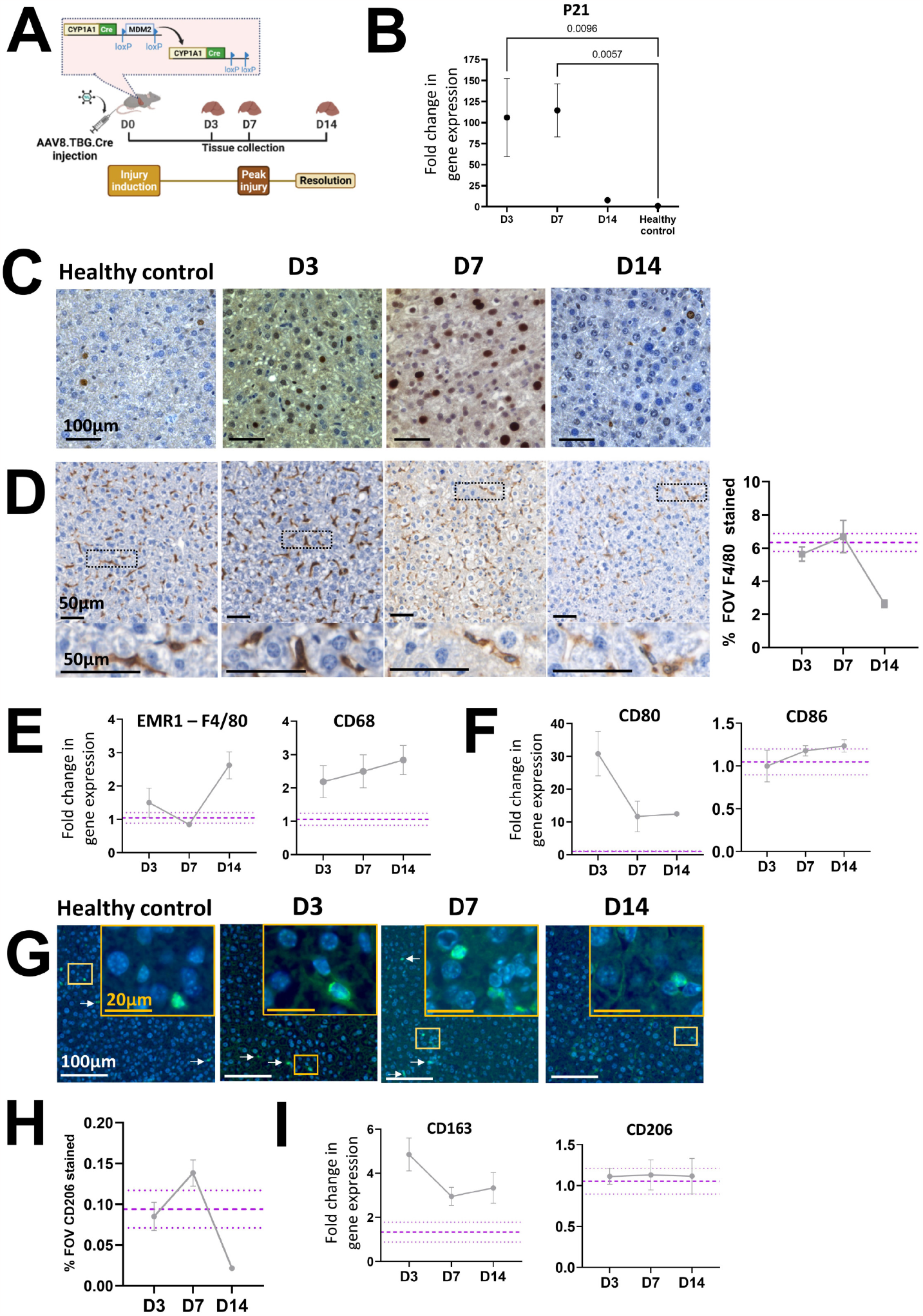
Induction of senescence in *Mdm2* mice by AAV8.TBG.Cre dosing leads to time sensitive inflammatory and regenerative macrophage responses. Purple dashed lines are healthy control mean±SEM. Data points represent mean of N=3-5 independent animals per timepoint analysed. All error bars are ±SEM. All qPCR **r**esults were normalised to the PPIA housekeeper and expressed as fold change relative to healthy control mean expression. For histological staining quantification the % of the field of view (FOV) positively stained was assessed, with ≥10 FOVs analysed per animal. **[A]** Experimental schematic for senescence dose response study. Inset demonstrates Cre recombinase expression mediated loss of *Mdm2* in hepatocytes as a result of injection of hepatotropic AAV8.TBG.Cre. Timepoints for tissue collection and analysis are indicated by days since injury induction (e.g. D3). **[B&C]** Expression of the p21 senescence marker at D3, D7 and D14 following AAV8.TBG.Cre induction analysed by qPCR and histological staining. Results show a significant, time sensitive increase in senescent marker expression, with expression resolving to healthy control animal levels by D14. Ordinary One-Way ANOVA relative to healthy control mean with Dunnett’s Multiple Comparisons test. **[D]** Representative histological micrographs and quantification of the pan macrophage marker F4/80 at D3, D7 and D14 post-induction, alongside healthy control. Scale bars 50μm. Results demonstrate no overall increase in the density of macrophages as a result of senescence driven injury and are supported by limited fold changes in gene expression of the pan macrophage markers EMR-1 (F4/80) and CD68, as analysed by qPCR **[E].[F]**Time sensitive changes in gene expression of the inflammatory macrophage markers CD80 and CD86 as a result of senescence induction. **[G]**Representative immunofluorescent micrographs of the pro-regenerative macrophage marker CD206 at D3, D7 and D14 post-induction, alongside healthy control. Scale bars 20μm and 100μm as indicated. White arrows indicate positively stained cells. Insets are digitally magnified 1.5x. . **[H]** Quantification of CD206 staining, showing a time sensitive increase in staining. **[I]** Analysis of gene expression of the pro-regenerative macrophage markers CD163 and CD206.

### Induction of senescence driven injury in Mdm2 mice by AAV8.TBG.Cre results in time sensitive inflammatory and regenerative macrophage responses

A range of immune cells, including natural killer cells, neutrophils, T-cells, and macrophages are activated in response to liver injury and factors secreted from senescent cells, contributing to the inflammatory response and subsequently repair^11^. For simplicity, here we focused on macrophage populations, which play a critical role in inflammation, regeneration and repair of liver injury^12,13^. Detailed macrophage classification is difficult to capture at the tissue level due to the large number of markers required to phenotype the full range of subtypes. Instead, we grouped macrophages into two key phenotypes, ‘M_1_’ pro-inflammatory and M_2_ ‘proregenerative’ cells.

Our experimental data showed limited changes in the total density (% field of view) of the macrophage population following senescence induction at this dose, relative to healthy control mice, based on mRNA expression and tissue staining of F4/80 (**Fig. 1D & E**). Analysis of mRNA expression of the pro-inflammatory (M_1_,CD80 and CD86) and pro-regenerative (M_2_, CD163 and CD206) macrophage markers revealed time-dependent changes in expression. CD80 expression is absent or expressed at very low levels on unstimulated monocytes and macrophages during homeostasis^14,15^, but expression is elevated in response to injury/inflammation. We saw this reflected in the *Mdm2* model where at D3 dramatically increased CD80 mRNA expression was seen relative to low homeostatic levels (**Fig. 1F**). Upregulation at D3 is indicative of pro-inflammatory stimulation, and expression decreased rapidly at D7 and D14, in support of a switch towards a pro-regenerative state at these timepoints. CD86 expression is typically upregulated faster than CD80, peaking within 6-24 hours^15,16^, therefore peak expression of CD86 may occur prior to D3 and was not captured here. These results suggested induction of a population level phenotypic switch towards M_2_ at D7, with decreased CD80 expression at D7 supported by a corresponding increase in expression of the M_2_ marker CD206 at the tissue level (**Fig. 1G &H**) and CD163 at the mRNA level (**Fig. 1I**). This supports our definition of ‘peak injury’ as D7, based on concurrent decreasing inflammatory marker expression and increasing regenerative marker expression.

### Senescence driven liver injury results in time sensitive changes in endothelial cell activation, myofibroblast activation and collagen-I deposition in the Mdm2 mouse model

Angiogenesis, stimulated as a result of the inflammatory microenvironment, is a common hallmark of chronic and acute liver injury^12,17^. Surprisingly, we found no increase in the number of endothelial cells following induction of senescence driven injury based on tissue quantification of ETS related gene (ERG) positive cells (**Supplementary Figure 3A&B**), presumably as a result of the acute nature of the induced injury. These results were supported at the gene expression level where no upregulation of classical endothelial markers such as CDH5 or VE-Cadherin was seen (**Supplementary Figure 3C**). However, we did see evidence of a phenotypic change in the endothelial cells following evaluation of expression of vascular cell adhesion molecule 1 (VCAM-1), intracellular adhesion molecule 1 (ICAM-1) and atypical chemokine receptor 3 (ACKR3), which are all predominantly expressed in endothelial cells and enable enhanced immune cell migration and liver regeneration. In support of other mouse and human liver injury^18^, an increased proportion of endothelial cells positive for VCAM-1 during the inflammatory phase of injury (D3 and D7) was demonstrated, which resolved to levels comparable to healthy controls by D14 (**Fig. 2A**), herein defined as ‘activated endothelial cells’. At the mRNA level only small fold changes in VCAM-1 expression were seen in the Mdm2 model relative to healthy mice, with comparable results seen for ICAM-1 and ACKR3 (**Fig. 2B**). Given that total liver tissue was analysed, and the vascular niche makes up only a small proportion of cells relative to the total liver parenchyma, it is perhaps unsurprising that we were unable to capture this change.

**Fig. 2.**
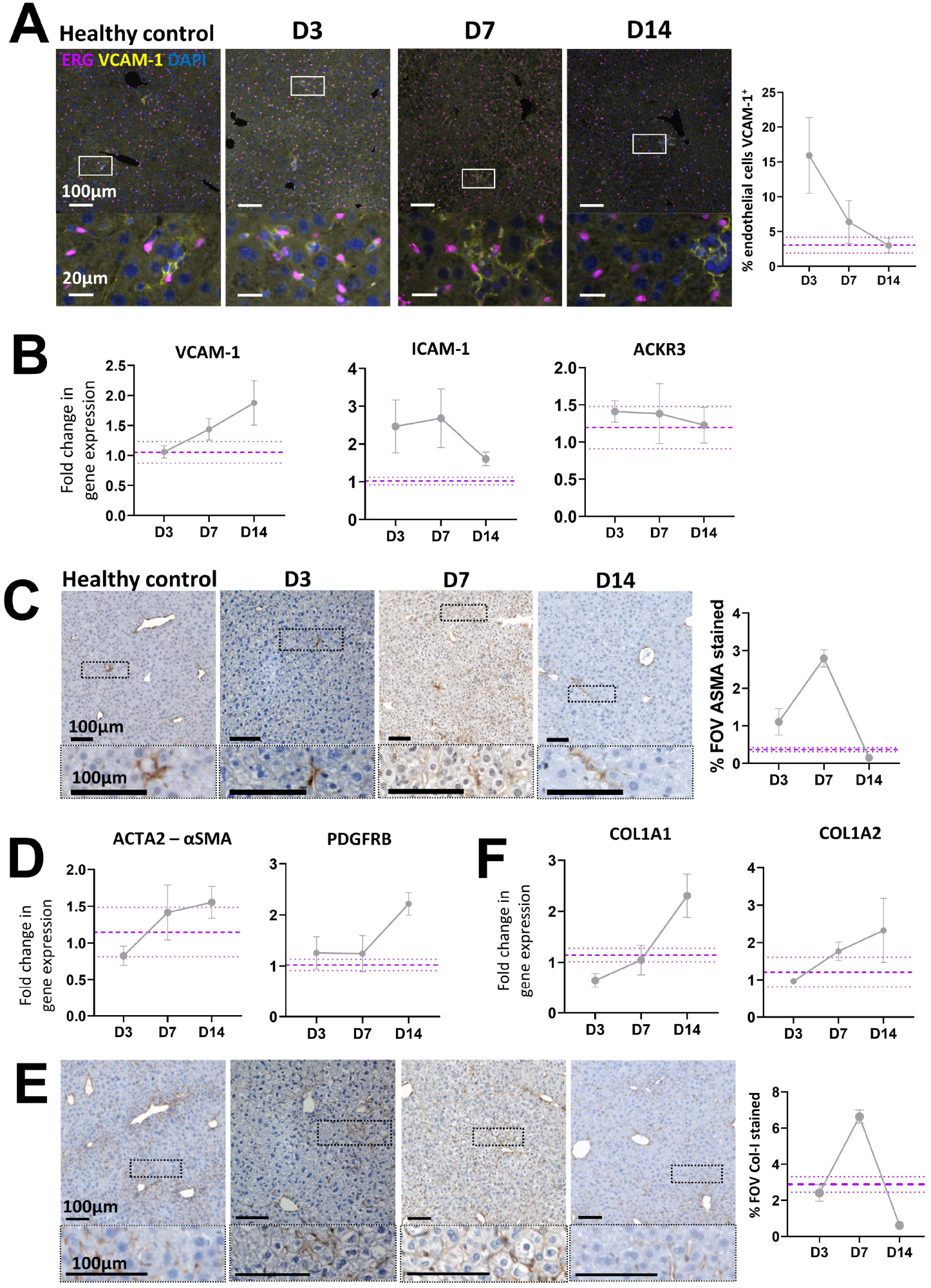
Induction of senescence driven injury in Mdm2 mice by AAV8.TBG.Cre results in time sensitive changes in endothelial cell activation, myofibroblast activation and collagen-I deposition. Purple dashed lines are healthy control mean±SEM. Data points represent mean of N=3-5 independent animals per timepoint analysed. All error bars are ±SEM. All qPCR results were normalised to the PPIA housekeeper and expressed as fold change relative to healthy control mean expression. For histological staining quantification of the % of the field of view (FOV) positively stained was assessed unless otherwise indicated, with ≥10 FOVs analysed per animal. **[A]**Representative immunofluorescent micrographs showing co-staining of the endothelial marker, ERG and activated endothelial cell marker VCAM-1 in healthy control animals and at D3, D7 and D14 following senescence induction, alongside quantification of ERG^+^VCAM-1^+^ cells. Results show a time sensitive increase in dual-positive endothelial cells indicating a time sensitive increase in this activated cell population, with expression resolving to healthy control animal levels by D14. **[B]**Fold change in total gene expression of the activated endothelial cell markers VCAM-1, ICAM-1 and ACKR3 following senescence induction. **[C]**Representative histological micrographs and quantification showing the time sensitive increase in staining for the activated myofibroblast marker α-SMA. Scale bars 100μm. **[D]**Analysis of changes in gene expression of the activated myofibroblast markers α-SMA and PDGFRB over time by qPCR. **[E]**Representative histological micrographs and quantification showing the time sensitive increase in staining for collagen-I. Scale bars 100μm. **[F]**Analysis of changes in gene expression of the pro-alpha1 and pro-alpha2 chains of collagen-I (COL1A1 and COL1A2) over time by qPCR.

Activated myofibroblasts, derived from a variety of cell sources, play an important role in scarring and wound healing in acute and fibrotic liver injury through tissue remodelling. Following liver regeneration and restoration of homeostasis, activated myofibroblasts are cleared as a result of apoptotic cell death or de-differentiation^19,20^. In the *Mdm2* model we see increased levels of α-SMA staining at D7, indicative of myofibroblast activation which reduces to within homeostatic levels by D14 as myofibroblasts clear (**Fig. 2C**). However, as with activated endothelial cells, at the mRNA level only subtle changes in expression of the α-SMA gene *Acta2* and the activated myofibroblast associated gene *Pdgfrb* (platelet derived growth receptor β isoform) were demonstrated (**Fig. 2D**). Again, this is likely a result of the relatively small contribution these cells make to the total liver cell population, thus capturing changes in mRNA expression of these markers from total liver is challenging. In support of myofibroblast activation, a corresponding increase in collagen-I deposition at peak injury is seen at the tissue level, with excess collagen-I resolved by D14 (**Fig. 2E**). Similarly to activated endothelial cells and myofibroblasts, limited changes in mRNA expression of the pro-alpha1 and pro-alpha2 chains of collagen-I relative to healthy controls were seen relative to those at the tissue level (**Fig. 2F**).

Overall, these results demonstrate that induction of senescence driven liver injury in the *Mdm2* model influences the key players in the liver microenvironment identified in other acute liver injury models. An initial inflammatory phase results in polarisation of macrophages towards an inflammatory state (**Fig. 1E**), facilitation of immune cell migration to the injury niche through activation of endothelial cells (**Fig. 2A**) and tissue remodelling as a result of myofibroblast activation (**Fig. 2C**) and collagen-I deposition (**Fig. 2E**). This is followed by a regenerative phase, including a macrophage phenotypic switch towards a pro-regenerative state (**Fig. 1F-H**), a reduction in activated endothelial cells and clearance of activated myofibroblasts and excess collagen-I (**Fig. 2A, C & E**).

### The ODE model accurately *captures* the senescence driven changes in the liver microenvironment

The variables and their interactions used in the mathematical model are shown in **Fig. 3A**. For more details about the ordinary differential equation model (ODE) development and the underlying assumptions we refer to **Materials & Methods**. Our results demonstrate that the theoretical predictions for the time-dependent evolution of each of the variables qualitatively capture the experimental observations (**Fig. 3B**), namely, the induced senescence decreases over time, returning to baseline as seen experimentally in **Fig1B&C**. All other populations (e.g. macrophages (**Fig. 1F**, CD80, pro-inflammatory and **Fig. 1G-I** pro-regenerative), the activated endothelial cells (**Fig. 2A**), activated myofibroblasts (**Fig. 2C**) and ECM (collagen-I (**Fig. 2E**)) initially increase before returning to baseline values (akin to healthy mouse levels). Having established that the mathematical model qualitatively captures the experimental data, we next used the model to determine how the system responds to differing levels of initial senescence. For the parameter sets considered, we found a critical threshold level of initial senescence which determines the transition between full resolution or irreversible injury due to uncontrollable inflammation.

**Fig. 3.**
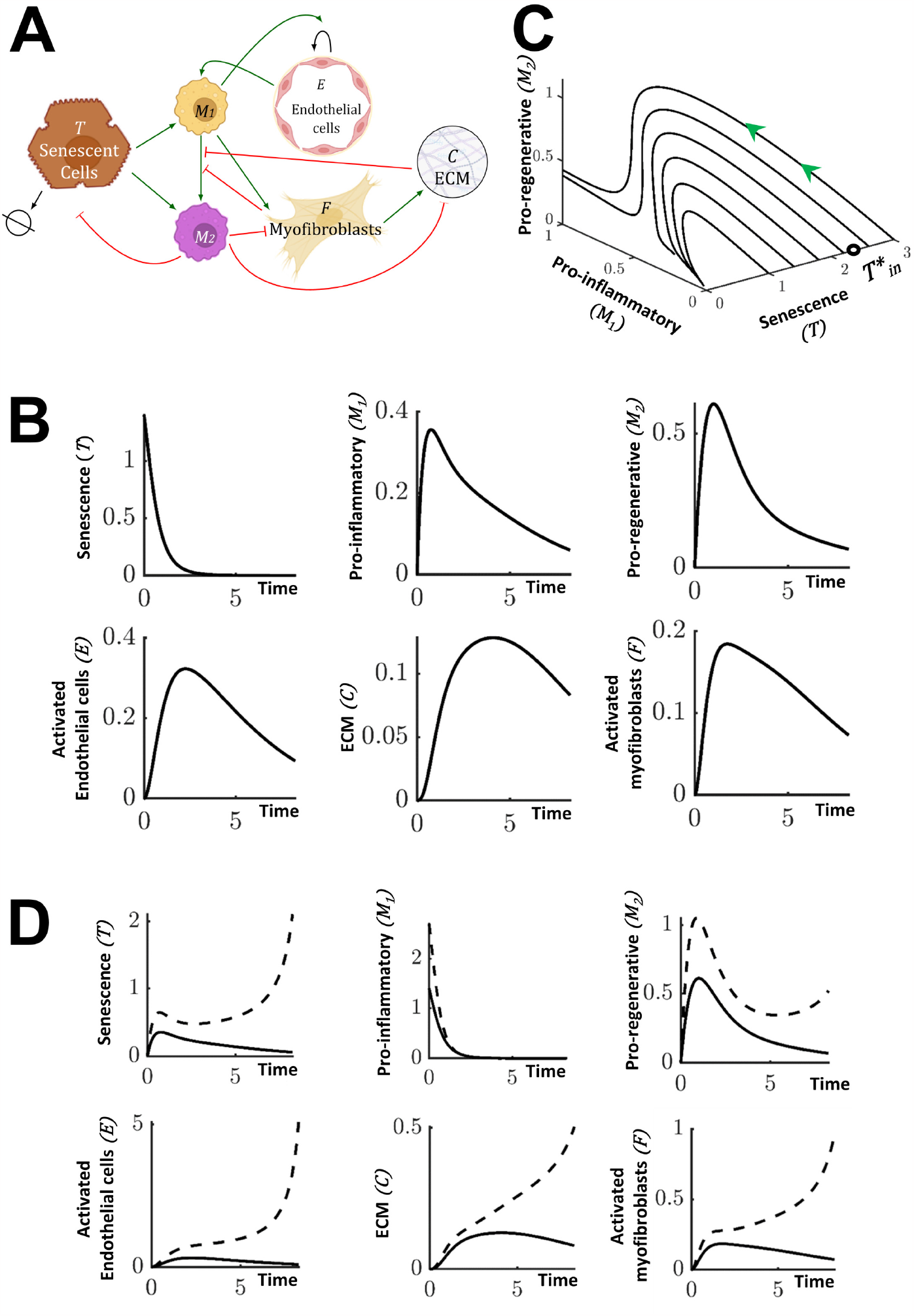
Mathematical model of senescence driven liver injury highlighting numerical predictions of the injury dynamics for a ‘medium’ senescence level. All numerical simulations were computed with parameter values set equal to 1, apart from the value of the initial senescence *T*_*in*_ which is given in the caption. **[A]**Schematic of the injury model where the variables and their interactions are presented. Green arrows indicate promotion and flat head red arrows indicate inhibition. **[B]**Mathematical prediction of the time evolution of all system components for a ‘medium’ initial senescence, *T*_*in*_ = 1.4. **[C]**Mathematical prediction of the evolution of M_1_ and M_2_ for different initial induced senescence levels. Above the critical initial senescence, denoted by 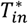 (black circle), the values of M_1_ and M_2_ do not return to the homeostatic values, indicating uncontrolled inflammation. **[D]**Mathematical prediction of the time evolution of all variables for the standard ‘medium’ *T*_*in*_ = 1.4 initial senescence (solid line) and at *T*_*in*_ = 2.7, which is above the critical initial senescence (dashed line). As in **[C]**, all the values do not reach a homeostatic level, but keep increasing.

We use the model to interrogate the influence of the initial senescence level on the key players in the liver injury microenvironment over time. **Fig. 3C** indicates how the pro-regenerative and pro-inflammatory macrophages evolve in time relative to the initial level of senescence. Each curve in **Fig. 3C** corresponds to a different level of initial senescence and models the evolution of the pro-inflammatory and pro-regenerative macrophage populations over time. We can see that the time evolution of the system depends on the initial senescence level. For all values of initial senescence, the level of senescence returns to its homeostatic value (**Fig 3D)**. If the initial senescence is less than the critical value 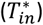, then the pro-inflammatory and pro-regenerative macrophage populations initially increase, before then returning to their homeostatic levels (**Fig. 3C**,**D)**. As the initial senescence level increases (but still below the critical initial senescence) the peak levels of pro-inflammatory and pro-regenerative macrophages increase (**Fig. 3C**). If we further increase the initial senescence levels to critical values exceeding 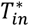 then the system will not resolve as the values of the variables (except senescence) do not return to zero, the homeostatic level. It is therefore crucial to identify and predict this critical initial senescence level 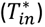, and determine how it depends on key system parameters (see below).

The explicit time dependence of each of the key players is revealed in **Fig. 3D** which indicates the response of the system when subject to moderate levels of initial senescence below the critical value (solid line) and levels of senescence above critical, resulting in uncontrollable inflammation (dashed line). Motivated by the *in vivo* data, we restrict attention to values of initial senescence below critical level as at these levels the inflammatory response will still be within the range which can ethically tested experimentally to validate the results of *in silico* predictions. Restricting attention to initial senescence levels below critical, **Fig. 3C** clearly indicates a dose response of the system, with higher levels of inflammation achieved prior to return to baseline for higher levels of initial senescence.

We now use the model to predict the system response to the induction of more severe (but still subcritical) and milder injury levels, and compare our model predictions with *in vivo* data.

### The ODE model accurately *predicts* microenvironmental changes following modification of initial levels of induced senescence

As demonstrated in **Fig. 3C**, our ODE model predicts that induction of higher and lower levels of initial senescence (restricted to below critical initial senescence) will result in a dose response of the system, with higher levels of inflammation achieved prior to return to baseline for higher levels of initial senescence. We experimentally tested the ODE predictions and performed a set of experiments across a range of AAV8.TBG.Cre doses and examined the resulting influence on the microenvironment at discrete timepoints. Seven distinct dosing groups were defined ranging from 5.0x10^9^ to 1.25x10^11^ GCU (*c*.*f*. materials and methods, groups 1-7), which our data from administering this dose range to Rosa26^LSL-TdTomato^ mice suggests induces recombination in 45%-96% of hepatocytes (**Supplementary Fig.1**). We focus here on the results of two additional doses 1.25x10^11^GCU (group 1, expected recombination >96%) and 5.0x10^9^GCU (group 7, expected recombination 45.39±12.935%) corresponding to induction of more severe and milder liver injury respectively than reported in **Fig. 1 and 2** (moderate injury, group 3) and report these in relation to the ODE predictions. Here, when referring to mathematically simulated levels of induced senescence we use the terms High, Medium and Low whereas biologically induced levels of senescence are referred to as severe, moderate (**Fig. 2**) and mild corresponding to the level of injury induced. These are not directly numerically linked as this study does not perform quantitative analysis, however formal calibration and validation will be able to achieve this in the future with the model suitably agile that when calibrated it can be applied to other systems where inflammation and fibrosis are key factors.

Firstly, we confirmed that doses for group 1 (severe injury) and group 7 (mild injury) resulted in comparatively higher and lower senescence induction, with the severe injury dose resulting in significantly increased p21 expression relative to healthy, age matched control mice at D3 (mean fold change 349.7±96.09) reaching comparable levels at D7 to a moderate dose (mean fold change 109.6±2.810, severe, and 114.6±31.53 moderate) before resolving by D14 (**Fig. 4A & B, Fig.1B**); whereas no significant increase in senescence was seen in group 7 relative to healthy controls. Results across all seven dosing groups also showed a linear trend of senescence induction based on senescent marker expression at D7 confirming the tight regulation of the level of induced senescence enabled by AAV8.TBG.Cre dosing in the *Mdm2* model (**Supplementary Figure 4**).

**Fig. 4.**
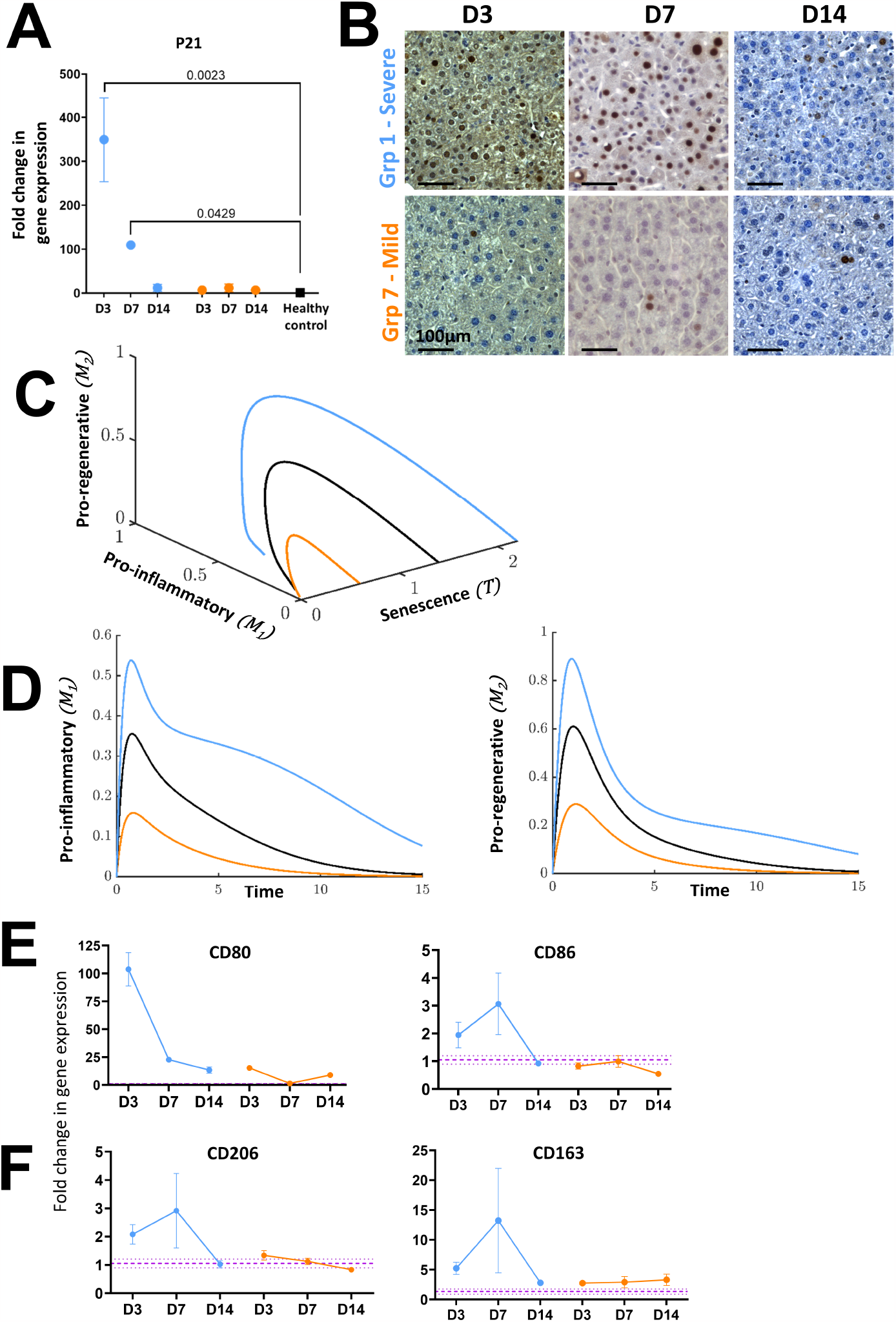
Mathematical predictions for the evolution of macrophage populations at High, Medium and Low doses of initial senescence match characterisation results following induction of severe and mild senescence driven liver injury in *Mdm2* mice. All numerical simulations were computed with parameter values set equal to 1, apart from the value of the initial senescence *T*_*in*_ which was varied as stated in captions. For numerical simulations, light blue, orange and black solid lines represent Low dose, High, and Medium initial senescence, *T*_*in*_ = 0.6, 2.2 and 1.4, respectively. For biological data, orange and light blue refer to severe (group 1) and mild (group 7) injury respectively. Purple dashed lines are healthy control mean±SEM. qPCR results were normalised to PPIA housekeeper and expressed as fold change relative to healthy control mean expression. N=3-5 mice per group and timepoint analysed. Error bars are SEM. **[A]**Group 1 and group 7 expression of p21 at D3, D7 and D14 following AAV8.TBG.Cre induction. Results show dose and time dependent changes in expression of this senescent marker. Ordinary One-Way Anova relative to healthy control mean with Dunnett’s Multiple Comparisons test. **[B]**Representative histological staining demonstrating dose and time dependent changes in hepatic p21 expression. **[C&D]** Mathematical prediction for evolution of the M1 and M2 macrophage populations for High, Medium, Low and critical initial senescence over time. Different levels of senescence induce a transient, senescence dependent increase in both macrophage populations before dropping to homeostatic values. **[E]**Change in gene expression of CD80 and CD86 M_1_ macrophage markers over time following severe and mild senescence driven injury induction in the *Mdm2* model. **[F]**Change in gene expression of CD163 and CD206 M_2_ macrophage markers over time following severe and mild senescence driven injury induction in the *Mdm2* model. Results in **[E]** and **[F]** demonstrate that the numerical predictions in **[C&D]** mirror the experimental trends.

The ODE model predicts a dose-dependent inflammatory and regenerative macrophage response, with proportional increases in pro-inflammatory and pro-regenerative macrophages determined by the level of initial senescence with all populations returning to baseline, homeostatic levels over time, provided the initial senescence is below the critical level (**Fig. 4C&D**). This prediction is in line with what is seen experimentally in the *Mdm2* model. As reported in **Fig. 1D** for moderate injury, there were limited changes in the total density of macrophages based on quantification of F4/80 staining relative to the induced level of senescence, this was consistent across the dose response (**Supplementary Figure 5A&B**). However, when considering more specific markers to identify the phenotype of the macrophage, our biological results qualitatively matched the ODE predictions. Induced senescence in group 1, corresponding to severe injury, resulted in dramatic upregulation in mRNA expression of the pro-inflammatory macrophage marker CD80 during the inflammatory phase (D3-D7), with residual upregulation of the earlier M1 marker CD86 also seen across this timeframe and expression of both markers reducing towards the levels of healthy control animals by D14, whereas only minor changes in CD80 were seen with mild injury induction (**Fig. 4E**). A comparable trend is seen for the M2 macrophage markers CD206 and CD163, where severe injury results in upregulated expression at D7 and downregulation following system resolution by D14, with mild injury showing comparable expression to healthy controls (**Fig. 4F, Supplementary Figure 5C&D**).

The ODE-model also predicts senescence dose-dependent changes in the activated endothelial cell population with proportional increases determined by the level of initial senescence and all populations returning to baseline over time (**Fig. 5A**). These predictions were confirmed experimentally. Induction of severe injury results in a transient increase in activated endothelial cells at the tissue level, and a corresponding transient increase in VCAM-1 mRNA expression, whereas mild injury results in expression comparable to healthy controls (**Fig. 5B-D**). During injury and inflammation VCAM-1 can also be expressed in other cells including Kupffer cells, hepatocytes and dendritic cells, this is demonstrated in the immunofluorescent micrographs (**Fig. 5B**) which show additional cells positive for VCAM-1 but negative for the endothelial marker ERG. In support of a phenotypic switch to activated endothelial status our results also demonstrated a dose dependent, transient increase in expression of the proinflammatory marker ICAM-1 and the pro-regenerative marker ACKR3 (**Fig. 5D**).

**Fig. 5.**
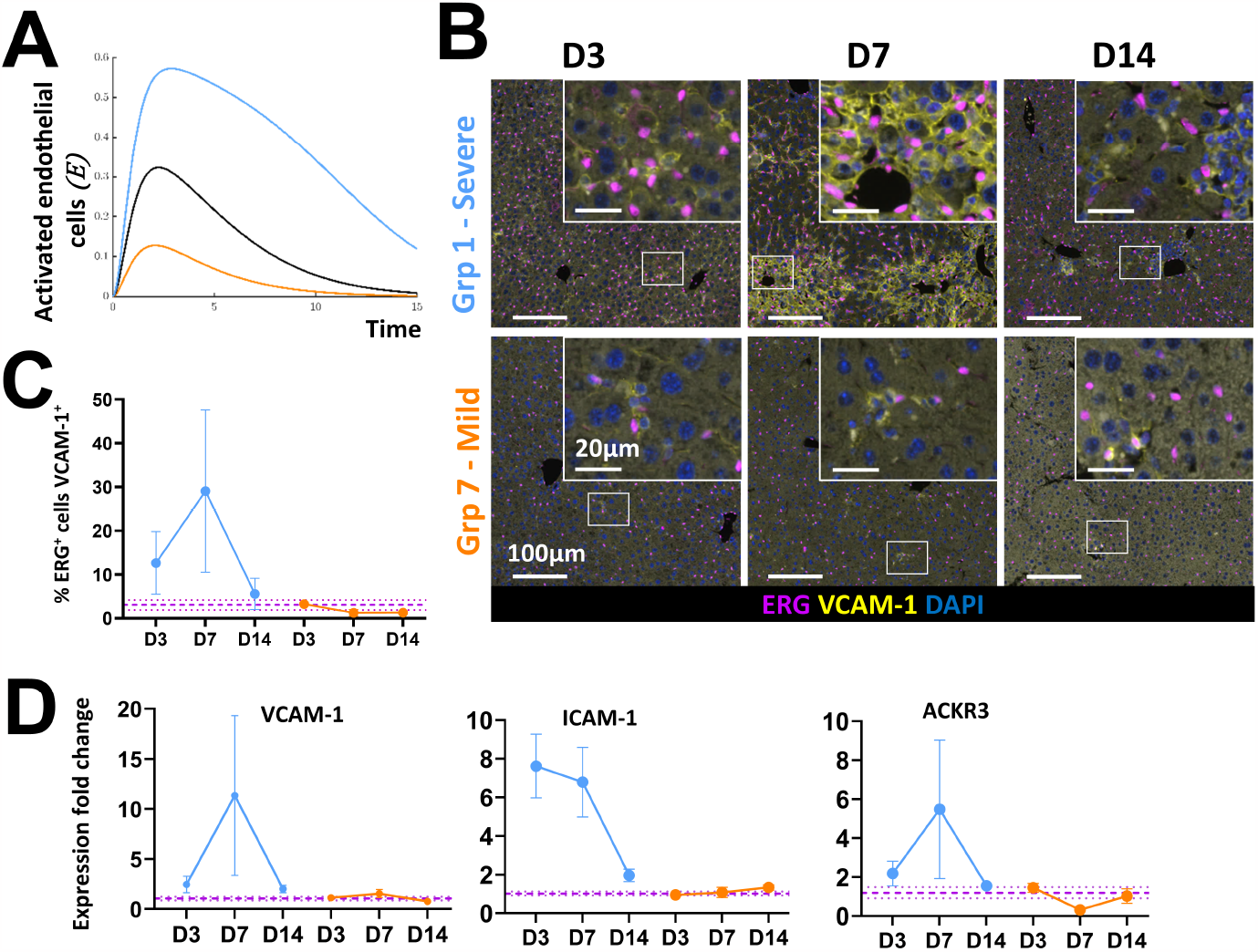
Mathematical predictions for the evolution of activated endothelial cells at High, Medium and Low doses of initial senescence match characterisation results following induction of severe and mild senescence driven liver injury in Mdm2 mice. All numerical simulations were computed with parameter values set equal to 1, apart from the value of the initial senescence*T*_*in*_ which was varied as stated in captions. For numerical simulations the light blue line represents the low dose, orange line the high and black the medium, *T*_*in*_ = 0.6, 2.2 and 1.4, respectively. For biological doses orange and light blue refer to severe (group 1) and mild (group 7) injury respectively. Purple dashed lines are healthy control mean±SEM. qPCR results were normalised to PPIA housekeeper and expressed as fold change relative to healthy control mean expression. N=3-5 mice per group and timepoint analysed. Error bars are SEM. **[A]**Mathematical prediction for evolution of activated endothelial cells for High, Medium, Low initial senescence. Different levels of senescence induce a transient, senescence dependent increase in activated endothelial cells before dropping to homeostatic values. **[B]**Representative immunofluorescent micrographs showing co-staining of the endothelial ERG marker and activated endothelial cell marker VCAM-1 at D3, D7 and D14 following senescence induction of severe and mild injury, alongside quantification of ERG^+^VCAM-1^+^ cells **[C]**. Results show a time and dose-dependent increase in dual-positive endothelial cells with expression resolving to healthy control animal levels by D14. **[D]**Fold change in total gene expression of the activated endothelial cell markers VCAM-1, ICAM-1 and ACKR3 following severe and mild senescence induction, supporting the immunofluorescence results. Numerical simulation results in **[A]** mirror these experimental trends.

Finally, the ODE-model also predicts senescence dose-dependent changes in the tissue microenvironment based on the temporal evolution of the activated myofibroblast cell population and ECM deposition. We depict the predictions for a High, Medium and Low level of initial senescence,*T*_*in*_ (**Fig. 6A&B**), which demonstrate increases in these populations relative to the level of initial senescence, with all populations returning to baseline over time (except where the critical initial senescence level is exceeded). These predictions accurately capture what is seen biologically when severe, mild (**Fig. 6C-H**) and moderate (**Fig. 2**) senescence driven injury are induced in the *Mdm2* model. We see peak activation of myofibroblasts at D7 based on α-SMA staining at the tissue level with more pronounced changes as a result of severe injury than mild injury and both dropping to healthy control levels by D14 (**Fig. 6C &D**). This is supported by upregulated mRNA expression of the α-SMA gene *Acta2* and *Pdgfrb* at D7 in severe injury (**Fig. 6E**). Similarly, we see peak and dose dependent deposition of collagen-I at D7 at both the tissue and mRNA level in severe and mild injury, which resolves to the level of healthy controls by D14 (**Fig. 6F & G**). This is supported by upregulated mRNA expression of *Col1a1* and *Col1a2* at D7 in severe injury (**Fig. 6H**). Comparable dose-dependent transient upregulation and downregulation at the tissue level for α-SMA and collagen-I was also seen across the full dose response (**Supplementary Figure 6**).

**Fig. 6.**
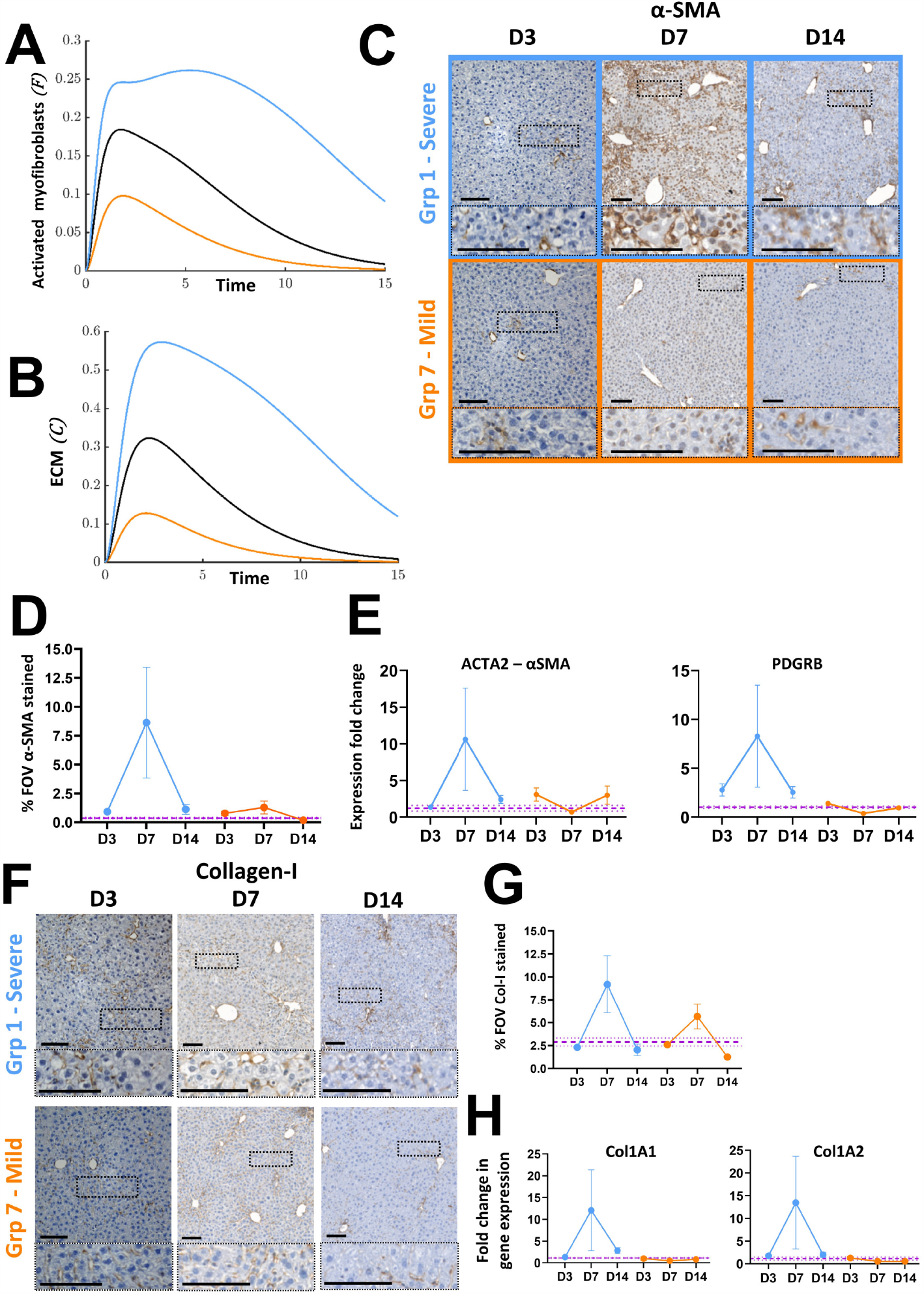
Mathematical predictions for tissue remodelling via evolution of activated myofibroblasts and ECM at High, Medium and Low doses of initial senescence match characterisation results following induction of severe and mild senescence driven liver injury in Mdm2 mice. All numerical simulations were computed with rates equal to 1. For numerical simulations the light blue line represents the low dose, orange line the high and black the medium level of senescence, *T*_*in*_ = 0.6, 2.2 and 1.4, respectively. For biological doses orange and light blue refer to severe (group 1) and mild (group 7) injury respectively. Purple dashed lines are healthy control mean±SEM. qPCR results were normalised to PPIA housekeeper and expressed as fold change relative to healthy control mean expression. N=3-5 mice per group and timepoint analysed. Error bars are SEM. Mathematical prediction for evolution of activated myofibroblasts **[A]** and ECM **[B]** for High, Medium and Low senescence over time. Different levels of senescence induce a transient, senescence dependent increase in both populations before dropping to homeostatic values. **[C&D]** Representative histological micrographs and quantification showing the transient, dosedependent increase in staining for the activated myofibroblast marker α-SMA. Scale bars 100μm. **[E]**Analysis of changes in gene expression of the activated myofibroblast markers α-SMA and PDGFRB over time by qPCR. **[F&G]** Representative histological micrographs and quantification showing the transient, dosedependent increase in staining for collagen-I. Scale bars 100μm. **[H]** Analysis of changes in gene expression of the pro-alpha1 and pro-alpha2 chains of collagen-I (COL1A1 and COL1A2) over time by qPCR.

### The ODE model predicts how different parameters impact the critical initial senescence level and senescent cell clearance time

Numerical simulations of the ODE model are much quicker and cheaper to perform than experiments, enabling rapid investigation of parameter space. Having demonstrated excellent qualitative agreement between the predictions of the theoretical model and the experimental data, we now exploit the model to determine the impact of system parameters on the critical initial senescence level, 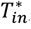, that can be tolerated by the system.

We predict how the rates of macrophage phenotype switch from pro-inflammatory to proregenerative (*G*) and increasing the activation rate of the endothelial population (*B*_*E*_) impact the critical initial senescence level. **Fig. 7A** shows how the critical initial senescence levels, 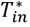, depend on *G* and *B*_*E*_. Note that we constrain parameters such that *G* > *B*_*E*_ − 1 (see *Materials & Methods – Mathematical model formulation*) to ensure resolution is always possible for subcritical initial senescence levels, and hence we do not consider the region of parameter space for which *G* < *B*_*E*_ − 1 (denoted “No resolution” in **Fig. 7A**). For a fixed rate of macrophage phenotype switch, increasing *B*_*E*_ decreases the critical initial senescence, as the pro-inflammatory nature of increasing B_E_ results in uncontrollable inflammation with lower initial senescence. Conversely, for a given rate of endothelial cell activation *B*_*E*_, increasing the macrophage phenotype switch (*G*) results in higher initial senescence levels being tolerated before uncontrolled inflammation is triggered. This is due to the proregenerative nature of this phenotypic switch, which will have a knock-on effect of reducing the activation of endothelial cells due to fewer *M*_1_ cells in the system.

**Fig. 7.**
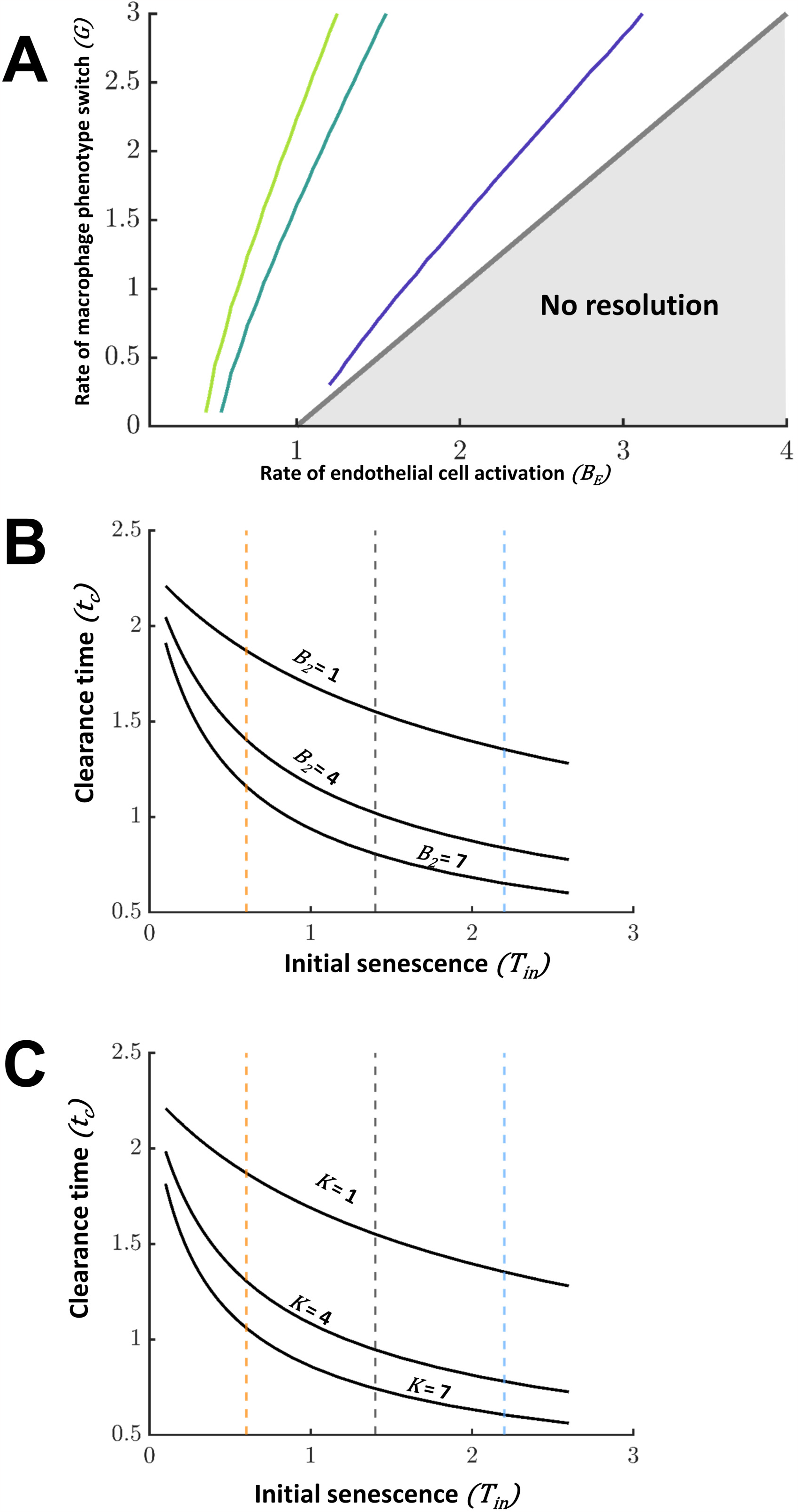
Mathematical predictions for critical initial senescence and clearance time based on the different rates of the system. Parameter study on the effect in critical initial senescence and clearance time (t_c_). **[A]**Effect of rate of macrophage phenotype switch (*G*) and endothelial cell activation *B*_*E*_ on critical initial senescence. All other parameters are set to 1. The pale green, dark green and purple lines are contours of 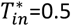, 5.5 and 10.5. An increase in *G* allows higher critical initial senescence while an increase in *B*_*E*_ permits smaller initial senescence levels before uncontrollable inflammation of the system is triggered. **[B]**Effect of the rate of *M*_2_ recruitment, *B*_2_, on clearance time for different initial levels of senescence, *T*_*in*_. An increase *B*_2_results in faster clearance and therefore lower t_c_. Orange, black and blue dashed lines represent Low, Medium and High initial senescence levels respectively. **[C]**Effect of the rate of senescent cell removal by *M*_2,_ *K*_*T*_, on clearance time for varying initial senescent values. An increase in this rate will result in faster clearance and therefore lower t_c_. Orange, black and blue dashed lines represent numerically Low, Medium and High initial senescence levels respectively.

The model can also be exploited to investigate the impact of parameters on the time for clearance of senescent cells within the system. The rate of removal of senescent hepatocytes

(*T)* depends on the number of *M*_2_ which in turn depends on the rate of their recruitment, *B*_2_, and the rate of *M*_2_-medicated removal of senescent ce11s, *K*. **Fig. 7B and C** predicts the effect of different initial senescence values on clearance time, which is defined as the time at which the senescent cell population reaches 10% of the initial value. Increased rates of *B*_2_ or *K* result in faster removal of the senescent population (and hence a lower t_c_). These simulations show that an enhancement of the parameters by modulating the micro-environment can result in lower injury levels. This highlights the potential use for mathematical models to aid development and predict outcomes in experimental interventional therapies.

## Discussion

The global incidence and prevalence of acute and chronic liver diseases is rising, despite advances in direct acting antivirals for hepatitis B and C. Non-alcoholic fatty liver disease is the most prevalent liver disease world-wide and alcohol related liver disease is on the rise in the UK. Outside of orthotopic liver transplantation, current therapies slow the progression of disease, but do not repair damaged tissue. Alternative therapies capable of promoting liver regeneration and restoring function are urgently required to support a growing patient burden. Alternative treatment approaches are being developed to try to tackle the clinical burden of severe liver disease. Regenerative medicine in the form of cellular therapy (macrophages, hepatocytes and stem cells) and tissue engineering (implantable constructs of hepatic tissue, bioartificial livers) approaches are of increasing interest in liver disease, but have not yet reached widespread clinical application. A number of hurdles need to be overcome to realise the clinical potential of regenerative therapies, including safety, efficacy and immunogenicity challenges. The successful clinical translation of regenerative medicine therapies requires robust preclinical tools. Complementary *in vivo* and *in silico* approaches can be used to advance pre-clinical studies.

Here, we developed an integrated *in silico-in vivo* model to capture the dynamics of key players in the regenerative response to acute senescence-mediated liver injury. We show that the mathematical model captures the host response to moderate liver injury via qualitative comparison of the model predictions with the *in vivo* experimental data. Importantly, we then show evidence of the predictive potential of *in silico* modelling, where we used our theoretical model to predict the host’s response to milder and more severe injury. These predictions were tested *in vivo*, where we obtained excellent qualitative agreement and validation of our mechanistic mathematical model. In addition, the model was able to determine a critical level of induced senescence that the system is unable to resolve due to uncontrolled inflammation. Whilst untested in this study for ethical reasons, the prediction fits with previous experimental experience of high dose AAV8.TBG.cre resulting in excess severity limits for mice. Of particular interest was how the numerical simulations could provide insight into potential mechanisms of therapy.

The depth of *in vivo* data included here did not always allow for experimental statistical significance, however, it provided an exciting opportunity to build a predictive mathematical model. The process of constructing the mathematical model was iterative, requiring decisions to be made about key players and their interactions. These were then refined through comparison of the mathematical model predictions and interrogation of the data. The experimental and theoretical variables selected were subjective and do not encompass the full complexity of the wound-healing response. Such a response involves numerous interactions between system components including multiple cell types (of which many are not captured in this study), the composition of the ECM and interstitial fluid across spatial and temporal scales^21^. Experimental data revealed a 30-fold increase in the expression of CD80 at the day 3 timepoint with moderate injury induction, this reached 100-fold in more severe injury, but only 10-fold with mild-induction. Given its role in T-cell activation, these data suggest exploration of the CD28 co-stimulatory pathway with possible incorporation of T cells into the mathematical model as a key parameter. Building and ensuring adequate model complexity with respect to the context of the biological question will improve the effectiveness of the mathematic approach.

Despite the necessary simplifications to obtain a qualitative framework, running numerical simulations enabled greater insight into potential mechanisms of therapy. By directly modulating different parameters in the system, we were able to assess how these different parameters impact the critical initial senescence level captured by the ODE model. For example, a faster macrophage phenotypic switch from pro-inflammatory to pro-regenerative, or the addition of pro-regenerative macrophages into the system, allowed for a higher level of initial senescence to be tolerated and for an accelerated rate of senescent cell clearance. These data indicate that targeted therapeutic strategies capable of increasing the rate of macrophage phenotypic switch in the liver could enhance regeneration in hepatic liver injury where high levels of senescence are evident. Alternatively, pre-conditioning treatment with adoptively transferred pro-regenerative macrophage therapy could also be beneficial. This hypothesis is in support of the use of macrophages for the treatment of liver cirrhosis^22^ (Phase II trial recently completed; ISRCTN10368050) and alternatively activated, pro-regenerative macrophages for experimental acetaminophen (APAP) liver injury^13^. Future studies can now focus on small experimental deviations but with higher n numbers for statistical power.

The ability to predict the host’s regenerative response following the adjustment of a key variable (in this case hepatocyte senescence) allows us to build on these predictions in the future with the addition of further experimental variables, such as intervention with a cellular therapy. Preclinical cellular transplantation studies may benefit substantially from the addition of predictive *in silico* modelling. We, and others, demonstrate the therapeutic potential of cellular therapies for liver disease, utilising multiple cell types (hepatocytes, hepatic progenitor cells, direct reprogrammed cells) in a range of experimental hepatocellular and biliary injury models^7,23–28^. A common feature uniting preclinical studies is significant variability in cell engraftment efficiency and therapeutic efficacy, which resonates with the variance observed in data from clinical trials^29^. Mathematical models can aid in standardisation of experimental data^8^. Our future studies aim to develop mathematical models that predict the effect of the host environment at the time of transplant on the outcome of cellular transplantation in preclinical models. Mechanistic modelling may be exploited to accelerate the translation of liver cell therapies into the clinic and opens up the prospect of developing personalised regenerative medicine.

## Materials and Methods

### Selection of key cellular players in the liver regenerative response

Senescent cells are metabolically active but non-proliferative and are capable of secreting a range of molecular factors (chemokines, cytokines, proteases and growth factors) collectively known as the senescence-associated secretory phenotype (SASP). SASP factors alter the cellular and structural composition of the surrounding microenvironment by inducing inflammation and immune cell recruitment, stimulating angiogenesis and modifying the extracellular matrix composition (e.g. through activation of myofibroblasts and subsequent upregulation of collagen synthesis^5,30^). Changes in these variables are reported in a variety of acute liver injury models^3,4,11^ and can be characterised at different stages of the injury and repair process in the *Mdm2* model through histological and gene expression analyses.

In this study we selected the key players to characterise in the experimental studies and interrogate in the mathematical model based on their previously reported critical roles in the inflammatory and/or regenerative phases of liver injury. A range of immune cells, including natural killer cells, neutrophils, T-cells, and macrophages are activated in response to liver injury and SASP factors, contributing to the initial inflammatory response and subsequently repair^11^. In this study we focused on macrophage populations, as they play a critical role across the entire repair process, contributing to both inflammation and regeneration of liver injury^12,13^. Pro-inflammatory macrophages (mathematically noted by *M*_1_) were considered to express classical macrophage markers such as CD80 and CD86. CD80 expression is absent or expressed at very low levels on unstimulated monocytes and macrophages during homeostasis^14,15^, but expression is elevated in response to injury/inflammation with highest expression notable on bone-marrow derived Kupffer cells recruited to sites of local inflammation^31^. Markers of alternative activation such as the scavenger receptors CD163 and CD206 were used to delineate pro-regenerative macrophages (M_2_ cells)^32^.

Macrophages support angiogenesis through co-localisation with newly formed vessels and subsequent secretion of cytokines, interleukins and growth factors which promote migration and proliferation of endothelial cells. Liver sinusoidal endothelial cells, as a core component of the hepatic vascular niche, secrete angiocrine factors such as hepatocyte growth factor and Wnt2 which have a paracrine effect stimulating liver regeneration^33,34^. Therefore, we also focused on investigating changes in the hepatic vascular niche across the injury repair process. As well as considering classical markers of endothelial cells that may reveal changes in vascular density/induction of angiogenesis, such as CD31 and vascular endothelial growth factor, we also investigated markers with differential expression by endothelial cells dependent on the stage of the injury repair process. Specifically, inflammatory cytokines such as TNF-A and IL-1B trigger vascular cell adhesion molecule 1 (VCAM-1) expression on the surface of endothelial cells which in turn induces adherence to and extravasation of leukocytes across blood vessel walls^35^. Increased endothelial expression of VCAM-1 therefore enhances immune cell infiltration to the site of injury. Similarly intracellular adhesion molecule 1 (ICAM-1), which is predominantly expressed in endothelial cells expression was also examined. ICAM-1 is also involved in the pro-inflammatory phase of injury by promoting leukocyte recruitment/transendothelial migration (T-cells, neutrophils, macrophages). Finally, atypical chemokine receptor 3 (ACKR3) is upregulated in response to acute liver injury where it acts as a receptor for CXCL12, functioning alongside CXCR4 to promote regeneration through secretion of angiocrine factors^36,37^.

Tissue remodelling is a key part of the injury repair process. Activated myofibroblasts play a pivotal and pleiotropic role in this process. Inflammatory cytokines such as IL-1β, TNF-α, CCL2, IL-6, transforming growth factor β1 (TGFβ1) and platelet derived growth factors (PDGFs) are produced by Kupffer cells and recruited macrophage populations following liver injury^38^. This inflammatory cascade results in activation of myofibroblasts within the injured microenvironment. Upon activation activated myofibroblasts express α-smooth muscle actin (α-SMA), matrix metalloproteinases (MMPs) and tissue inhibitors of metalloproteinases (TIMPs) as well as synthesize ECM proteins, including Collagen type I, the main component of fibrous scar^39^. Platelet derived growth factor receptor β isoform (PDGFRβ) acts as the primary PDGFR isoform to mediate activation and profibrogenic transdifferentiation of hepatic stellate cells into myofibroblasts during hepatic fibrosis^40^ and therefore this was used as an additional mRNA expression marker to assess changes in the activated myofibroblast population over time. We note that activated myofibroblasts can also be derived from other cell sources, such as portal fibroblasts and BM-derived fibrocytes, and these will also be captured by the assessment of their shared characteristics (changes in α-SMA and ACTA2 expression). Scar resolution is associated with recruitment and repolarisation of macrophages towards a pro-regenerative phenotype after the initial inflammatory cascade, wherein they secrete IL-10, MMPs and collagenases, inhibit the expression of TIMPs and phagocytose ECM fragments^32,38,41^. Activated myofibroblasts are subsequently cleared from the niche as a result of apoptotic cell death or de-differentiation/inactivation into a state where they cease collagen production^19,20^

### Iterative cycle

An iterative cycle of predict-test-predict-refine underpinned the development of the mathematical model. Initial theoretical model development was guided by existing literature and understanding of the host response to liver injury (see *‘Selection of key cellular players’*), allowing hypotheses to be made for the causal mechanisms underlying the biological system, which were then represented mathematically. Model predictions were then compared with the experimental data, and the mathematical models were refined as necessary. Once qualitative agreement was obtained, the mathematical model was exploited to predict the response of the host to mild and severe injury.

### Animals

Both male and female AhCreMdm2^fl/fl^ mice *(Mdm2)*^7,42^ and Rosa26^LSL-TdTomato^ mice on a C57Bl/6 background were used in this study. Genotyping was performed by the Transnetyx genotyping service. Animals were all within 14-27 weeks old at the start of the experiments. Mice were maintained in pathogen-free conditions with sterile food and water available *ad libitum* and kept in individually ventilated plastic cages with environmental enrichment and bedding material. Cages were held in dedicated, licenced air-conditioned animal rooms, under light/dark cycles lasting 12 hours daily. Maximum cage occupancy was six animals and fresh cages were supplied at least weekly. All animal experiments were carried out under procedural guidelines, severity protocols and with ethical permission from the University of Edinburgh Animal Welfare and Ethical Review Body (AWERB) and the Home Office (UK). Power calculations were not routinely performed, but animal numbers were chosen based on the anticipated magnitude of response from researchers’ experience with these strains.

AAV8.TBG.Cre virus at the concentrations indicated was diluted in sterile PBS to a final volume of 100μl and administered by tail vein injection to induce recombination. Dosing groups were randomly assigned with the following virus concentrations administered (GCU): 1.25x10^11^ (Group 1, “Severe” dose), 6.25x10^10^ (Group 2), 4.16x10^10^ (Group 3, “Moderate” dose), 3.13x10^10^ (Group 4), 2.50x10^10^ (Group 5), 1.25x10^10^ (Group 6), 5.00x10^9^ (Group 7, “Mild” dose) plus 4.17x10^9^ and 2.50x10^9^ (Rosa26^LSL-TdTomato^ mice only, *cf*. **Supplementary Fig.1**). Experimental endpoints were 3 days (Severe, Moderate and Mild dose groups, *Mdm2* mice), 7 days (all dosing groups, *Mdm2* mice) or 14 days (all dosing groups both *Mdm2* and Rosa26^LSL-TdTomato^ mice) after recombination induction with N=3-5 animals per group and timepoint analysed. Mice were euthanized according to UK Home Office regulations. Blood was collected by cardiac puncture and livers were perfused with PBS via the inferior vena cava to clear remaining blood. Subsequently, organs were harvested and either snap frozen directly at -80°C or fixed in 10% formalin (in PBS) for 8 hours and stored in 70% EtOH prior to embedding in paraffin blocks. Animals reaching experimental severity protocol boundaries were excluded from analysis, otherwise all animals were included.

### Immunohistochemistry

4μm thick sections of formalin-fixed paraffin embedded tissue blocks were collected onto glass slides, dewaxed and rehydrated prior to antigen retrieval. For DAB-stained sections, tissue was blocked with Bloxall (Vector), Avidin/Biotin block (Invitrogen), and protein block (Spring Bioscience). Sections were subsequently incubated at 4°C overnight with primary antibodies (concentrations and antigen retrieval conditions detailed in **Supplementary Table 1**). Primary antibody detection was via incubation with species-specific secondary biotinylated antibodies for 30 minutes at room temperature (Vector) followed by R.T.U. Vectastain (30 mins, room temperature), ABC reagent (Vector), and DAB chromogen (Dako). Slides were counterstained with haematoxylin. For immunofluorescence, sections were blocked for 30 minutes in protein block, incubated overnight with primary antibodies, washed extensively with PBS and incubated with Alexa 488 and Alexa647 conjugated secondary antibodies (Invitrogen) for 30 minutes at room temperature. Sections were counter-stained with DAPI (4′,6-diamidino-2-phenylindole) and mounted with Fluoromount-G (SouthernBiotech).

### Immunohistochemistry quantification

Immunostained slides were imaged on a Vectra Polaris (Perkin Elmer) or a Nikon Eclipse Ni microscope using a Teledyne Q-Imaging MicroPublisher 6 camera at 20 or 40x magnification. At least 10 randomly selected fields of view were imaged per animal. Quantification of the proportion of each field of view positively stained was performed using inForm software (CD206, immunofluorescence, PerkinElmer), or using macros developed with Fiji software (ImageJ) where intensity thresholds were set based on the isotype control for each marker, which was stained in parallel (**Supplementary Methods**). For VCAM-1 and ERG co-staining double positive cells were quantified using a pipeline developed in inForm software.

### RNA extraction and cDNA synthesis

Total RNA was extracted from 30-50mg samples of snap frozen liver tissue. Tissue was homogenised in RLT buffer. Subsequently RNA was extracted from tissue lysates using the RNeasy MiniKit (Qiagen) according to manufacturer’s instructions. RNA concentration and purity of isolated RNA was determined with a NanoDrop spectrophotometer. Samples with a 260/280 ratio ≥1.8, indicative of good quality RNA, were used for complementary DNA synthesis (cDNA). CDNA was synthesised by reverse transcription using the Quantitect Reverse Transcription Kit (Qiagen) according to manufacturer’s instructions. Any contaminating genomic DNA was removed with gDNA Wipeout Buffer.

### Quantitative real-time PCR

Analysis of mRNA expression of marker genes for the parameters of interest was performed by qPCR using the Fast SYBR green Master Mix (Thermofisher) on a Roche Lightcycler 480 using 12.5ng of cDNA per reaction. Cycling conditions were: pre-incubation (95°C, 20 seconds, 4.8°C/s), Annealing/Extension (95°C for 3s at 4.8°C ramp rate, 60°C for 30 seconds at 2.5°C/s ramp rate, for 40 cycles), Cooling (95°C, 30 seconds at 2.5°C/s ramp rate). Triplicate technical replicates for each biological sample were assayed for each gene. Gene specific primers used are detailed in **Supplementary Table 2**. Sample threshold cycle (C^t^) values were normalised to the murine peptidylprolyl isomerase (PPIA) housekeeping gene (ΔCt = meanC_t_^target gene^-meanC_t_^housekeeping gene^). For each gene the mean ΔCt for the healthy animal group was calculated (N=5 biological replicates) and set as the experimental control. The 2^*-ΔΔCt*^ method was subsequently used for analysis^19^, with results for each animal expressed as the fold change in expression relative to the healthy control average. Data points in gene expression graphs represent the mean fold change in expression for all biological replicates in each dosing group±SEM (N=3-5 animals per group).

### Mathematical model formulation

Variables *T, M*_l_, *M*_2_, *E*, and *F* describe the number of senescent cells, pro-inflammatory macrophages, pro-regenerative macrophages, activated endothelial cells and activated myofibroblasts, respectively. The ECM concentration is denoted by C. In a similar manner to the experimental model, we do not distinguish between primary or ‘secondary’ senescent cells, and instead consider a single population of senescent cells *T*. All cell populations and ECM concentration correspond to dynamic fluctuations from homeostasis, which arise due to the addition of senescent cells. The key interactions between the variables are illustrated in **Fig. 3A**. The senescent cells, macrophage populations and activated endothelial cell populations are normalised with respect to the homeostatic endothelial cell population, the myofibroblasts with respect to the homeostatic myofibroblast populations and the ECM concentration with respect to the homeostatic ECM concentration, so that dynamic changes in the variables represent fold changes for homeostatic levels. Time is denoted by t, normalised on the timescale for recruitment of macrophages by the activated endothelial cells.

The mathematical model abstracts the interactions (see *‘Selection of key cellular players’*) into ordinary differential equations using the law of mass action. The equations are as follows:

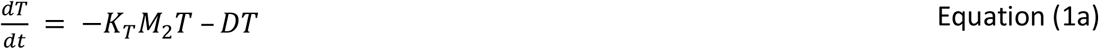

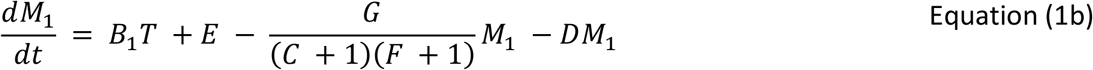

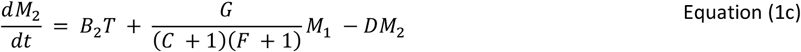

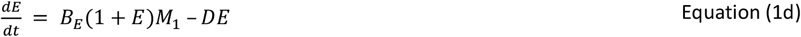

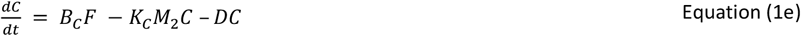

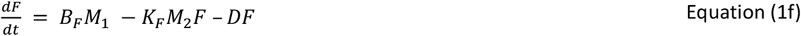

The terms on the left hand side (LHS) of equations (1a)-(1f) denote the rate of change of variables with respect to time. The assumption underpinning the terms on the right hand side (RHS) of equations (1a)-(1f) are as follows. The final term on the RHS of each of the equations (1a)–(1f) captures the loss of the variable of interest due to death or deactivation, with the parameter D capturing the rate of death/deactivation, e.g. in equation (1f) the term −DF captures the apoptotic cell death or de-differentiation/inactivation of the activated myofibroblasts. The pro-regenerative *M*_*2*_ population inhibits the generation of secondary senescent cells via secreted anti-inflammatory cytokines and chemokines (e.g. IL-10) and removes the primary senescent cells via phagocytosis. The resulting reduction in the total senescent cell population is modelled via the term −K_*T*_*M*_2_*T* in equation (1a) where *K*_*T*_ captures the rate of removal of senescent cells per pro-regenerative macrophage. The first term on the RHS of equations (1b) and (1c) models the recruitment of pro-inflammatory and pro-regenerative macrophages to the injured liver by the senescent cells at rates *B*_1_ and *B*_2_ per senescent cell respectively. The second term on the RHS of equation (1b) captures the recruitment of pro-inflammatory macrophages due to the activated endothelial cells. Note that no constant rate parameter appears in this term, reflecting the normalisation of time on the timescale for the recruitment of macrophages by the activated endothelial cells. Changes in the local environment impact macrophage polarisation with *M*_1_ switching to an *M*_2_ phenotype. The rate of this transition is impacted by ECM and activated myofibroblast generation, under the influence of pro-inflammatory (e.g. IFN-γ, TNF-⍰) and anti-inflammatory cytokines (e.g. IL-10)^10^. Excess ECM deposition results in an increased tissue stiffness whichdrives further recruitment of infiltrating immune cells and myofibroblast activation^43,44^. Accumulated activated myofibroblasts secrete IL-6, TIMPs, TGF-β, vascular endothelial growth factor (VEGF), epidermal growth factor (EGF), and CXCL10 which can reduce the polarisation of macrophages towards an *M*_2_ phenotype, impacting scar resolution and ECM turnover^32,38,41^. Turning to equation (1d), the *B*_*E*_*M*_1_ term represents activation of the homeostatic endothelial cell population due to the inflammatory environment during injury, which we assume depends on the concentration of pro-inflammatory macrophages, for simplicity. Here *B*_*E*_ is the rate of endothelial cell activation per pro-inflammatory macrophage. *The B*_*E*_ *E M*_1_ represents proliferation of the activated endothelial cell population in response to local inflammation. In equation (1e), *B*_C_F corresponds to ECM deposition by the myofibroblasts^10^ with *B*_C_ the deposition rate per myofibroblast, and *K*_C_ M_2_ C denotes breakdown of the ECM by proregenerative macrophages, where *K*_C_ is the rate of breakdown per pro-regenerative macrophage^10^. Finally, we consider equation (1f). Secretion of growth factors, e.g. TGF-⍰, by the pro-inflammatory macrophages stimulates differentiation of fibroblasts into contractile myofibroblasts which we model via the term *B*_*F*_*M*_1_ where *B*_*F*_ is the rate of differentiation per pro-inflammatory macrophage. The second term captures deactivation of activated myofibroblasts with *K*_F_ the deactivation rate per pro-regenerative macrophage.

Equations (1a-d) are solved subject to the following initial conditions

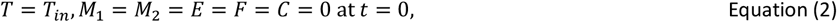

where *T*_*in*_ captures the initial senescent cell population as a result of the administration of the AAV8.TBG.Cre virus, and all other variables are set to zero indicating the populations are initially at their homeostatic levels.

The mathematical model has eleven parameters: *K*_*T*_, *D, B*_*l*_, *B*_2_, *G, B*_*E*_, *B*_*C*_, *K*_*C*_, *B*_*F*_, *K*_*F*_ and *T*_*in*_, each corresponding to an underlying mechanism. The scope of this paper is to determine the impact of key parameters on the system dynamics, which will allow us to determine the dominant mechanisms in acute senescence driven injury and repair. It is beyond the scope of this study to undertake a comprehensive parameter sweep, and instead we focus on five key parameters of interest: *T*_*in*_, *B*_*E*_, *G, B*_2_ and *K*_*T*_, and set all other parameter values to be 1 throughout this paper. Varying *T*_*in*_ models the different levels of induced senescence in response to administration of the AAV8.TBG.Cre virus. The long-time steady states admitted by the system do not depend on *B*_2_ and *K*_*T*_ (as from equation (1a) we see that the steady solution has *T* = 0 and hence parameters multiplying *T* do not feature in the steady state solution). We explore the impact of *B*_*E*_, the rate of endothelial cell activation per proinflammatory macrophage, and *G*, the rate of phenotypic switching from pro-inflammatory to pro-regenerative macrophages, as these processes underpin the competition between inflammation and regeneration, and (together with the death rate D) determine whether the system admits one or two steady states (see Supplementary Material). We further explore the impact of *B*_2_, the rate of pro-regenerative macrophage recruitment, and *K*_*T*_, the rate of clearance of senescent cells per pro-regenerative macrophage, on the system dynamics, acknowledging that the strength of the model is that it facilitates exploration of all parameter values.

Steady-state analysis of the governing equations reveals that two possible steady states exist, depending on parameter values: the trivial steady state where all variables are zero (corresponding to the system returning to homeostasis) and the non-zero steady state where the senescent cell population is zero but all other variables take non-zero positive values.

When *G* < *B*_*E*_ − 1 only a single steady state exists which is linear unstable, so that in practice this state can never be reached (small perturbations from the zero steady state will grow), resulting to uncontrollable inflammation. When *G* > *B*_*E*_ − 1 the system admits both steady states, with the trivial steady state being linearly stable while the non-trivial steady state is linearly unstable. Motivated by the experimental data which demonstrates that the system resolves to homeostatic values for the selected doses of the AAV8.TBG.Cre virus, we constrain the model to parameter set such *G* > *B*_*E*_ − 1. For details of the steady states and linear stability analysis, see Supplementary Material.

The ODE system is solved in Matlab using *ode45*. We confirmed model convergence by refining the time step and ensuring the solution did not change. For the steady states and the stability analysis, the variables are set as symbolic objects. We then find the Jacobian of the system using the *jacobian* function and finally we solve the steady state system using *solve*.

## ACKNOWLEDGEMENTS

The authors thank the University of Edinburgh Institute for Regeneration & Repair Imaging core (notably M. Vermeren and J. Cholewa-Waclaw) for imaging support and analysis assistance, and W. Mungall and M.Struzik for help with animal experiments. Elements of Figures in this paper were produced using biorender.com

## AUTHOR CONTRIBUTIONS (CRediT statement)

Conceptualization: *CAH, EA, WYL, SJF, VLG, SLW;* Collaborative paper structure: *CAH, EA, SMF, WYL, SJF, VLG, SLW;* methodology: Experimental design - *VLG, CAH*; Mathematical model development, analysis and solution - *EA, SLW*; software: *CAH, EA*; validation: *CAH, VLG, EA, SLW*; formal analysis: *VLG, CAH, EA, SLW, SJF*; investigation: *CAH, EA, RA, TYM, AS, VLG*; resources: *VLG, SJF, SLW*; data curation: *CAH, VLG, EA, SLW*; writing—original draft preparation: *CAH, EA, VLG, SLW*; writing—review and editing: all authors; visualization: *CAH, EA, VLG, SLW*; supervision: *WYL, SJF, VLG, SLW*; project administration: *CAH, EA, WYL, SJF, VLG, SLW*; funding acquisition: *WYL, SJF, VLG, SLW*. All authors have read and agreed to the published version of the manuscript.

## ETHICS STATEMENT

The study involved animals. The study was conducted in accordance with the Declaration of Helsinki. All procedures related to animal work were performed in accordance with all legal, ethical, and institutional requirements (PPL numbers 70/7847 and P231C5F81) as dictated by UK legislation. Sample sizes of *in vivo* experiments were based on prior work from our laboratory and kept to a minimum in line with law and ethical guidelines for animal research in the UK as were the *in vivo* endpoints.

## DATA AVAILABILITY STATEMENT

The data presented here are available on request from the first or corresponding author(s).

## References

1. Marcellin, P. & Kutala, B. K. Liver diseases: A major, neglected global public health problem requiring urgent actions and large-scale screening. Liver Int. 38, 2–6 (2018).

2. Bird, T. G. et al. TGFβ inhibition restores a regenerative response in acute liver injury by suppressing paracrine senescence. Sci. Transl. Med. 10, 1–15 (2018).

3. Ferreira-Gonzalez, S. et al. Paracrine cellular senescence exacerbates biliary injury and impairs regeneration. Nat. Commun. 9, 1–15 (2018).

4. Ogrodnik, M. et al. Cellular senescence drives age-dependent hepatic steatosis. Nat. Commun. 8, (2017).

5. Aravinthan, A. D. & Alexander, G. J. M. Senescence in chronic liver disease: Is the future in aging? J. Hepatol. 65, 825–834 (2016).

6. Aravinthan, A. et al. Hepatocyte senescence predicts progression in non-alcohol-related fatty liver disease. J. Hepatol. 58, 549–556 (2013).

7. Lu, W.-Y. et al. Hepatic progenitor cells of biliary origin with liver repopulation capacity. Nat. Cell Biol. 17, 973–983 (2015).

8. Waters, S. L., Schumacher, L. J. & El Haj, A. J. Regenerative medicine meets mathematical modelling: developing symbiotic relationships. npj Regen. Med. 6, 1–8 (2021).

9. Waters, S. L., Schumacher, L. J. & El Haj, A. J. Regenerative medicine meets mathematical modelling: developing symbiotic relationships. npj Regen. Med. 6, (2021).

10. Friedman, A. & Hao, W. Mathematical modeling of liver fibrosis. Math. Biosci. Eng. 14, 143–164 (2017).

11. Koyama, Y. & Brenner, D. A. Liver inflammation and fibrosis. J. Clin. Invest. 127, 55–64 (2017).

12. You, Q. et al. Role of hepatic resident and infiltrating macrophages in liver repair after acute injury. Biochem. Pharmacol. 86, 836–843 (2013).

13. Starkey Lewis, P. et al. Alternatively activated macrophages promote resolution of necrosis following acute liver injury. J. Hepatol. 737, 349–360 (2020).

14. Freeman, G. J. et al. Structure, expression, and T cell costimulatory activity of the murine homologue of the human B lymphocyte activation antigen B7. J. Exp. Med. 174, 625–631 (1991).

15. Fleischer, J. et al. Differential expression and function of CD80 (B7-1) and CD86 (B7-2) on human peripheral blood monocytes. Immunology 89, 592–598 (1996).

16. Lenschow, D. J. et al. Expression and functional significance of an additional ligand for CTLA-4. Proc. Natl. Acad. Sci. U. S. A. 90, 11054–11058 (1993).

17. Fernández, M. et al. Angiogenesis in liver disease. J. Hepatol. 50, 604–620 (2009).

18. Pirisi, M. et al. Serum Soluble Vascular-Cell Adhesion Molecule-1 (Vcam-1) in Patients with Acute and Chronic Liver Diseases. Dis. Markers 13, 11–17 (1996).

19. Desmouliere, A., Redard, M., Darby, I. & Gabbiani, G. Apoptosis mediates the decrease in cellularity during the transition between granulation tissue and scar. Am. J. Pathol. 146, 56–66 (1995).

20. Kisseleva, T. et al. Myofibroblasts revert to an inactive phenotype during regression of liver fibrosis. Proc. Natl. Acad. Sci. U. S. A. 109, 9448–9453 (2012).

21. Pellicoro, A., Ramachandran, P., Iredale, J. P. & Fallowfield, J. A. Liver fibrosis and repair: Immune regulation of wound healing in a solid organ. Nat. Rev. Immunol. 14, 181–194 (2014).

22. Brennan, P. N. et al. Study protocol: a multicentre, open-label, parallel-group, phase 2, randomised controlled trial of autologous macrophage therapy for liver cirrhosis (MATCH). BMJ Open 11, e053190 (2021).

23. Hallett, J. M. et al. Human biliary epithelial cells from discarded donor livers rescue bile duct structure and function in a mouse model of biliary disease. Cell Stem Cell 29, 355-371.e10 (2022).

24. Sampaziotis, F. et al. Cholangiocyte organoids can repair bile ducts after transplantation in the human liver. Science (80-.). 371, 839–846 (2021).

25. Azuma, H. et al. Robust expansion of human hepatocytes in Fah-/-/Rag2 -/-/Il2rg-/-mice. Nat. Biotechnol. 25, 903–910 (2007).

26. Overturf, K., Al-Dhalimy, M., Finegold, M. & Grompe, M. The repopulation potential of hepatocyte populations differing in size and prior mitotic expansion. Am. J. Pathol. 155, 2135–2143 (1999).

27. Wang, X. et al. The origin and liver repopulating capacity of murine oval cells. Proc. Natl. Acad. Sci. U. S. A. 100, 11881–11888 (2003).

28. Katsuda, T. et al. Conversion of Terminally Committed Hepatocytes to Culturable Bipotent Progenitor Cells with Regenerative Capacity. Cell Stem Cell 20, 41–55 (2017).

29. Hansel, M. C. et al. The history and use of human hepatocytes for the treatment of liver diseases: The first 100 patients. Curr. Protoc. Toxicol. 14.12.1-14.12.23 (2014) doi:10.1002/0471140856.tx1412s62.

30. Elder, S. S. & Emmerson, E. Senescent cells and macrophages: Key players for regeneration? Open Biol. 10, (2020).

31. Klein, I. et al. Kupffer cell heterogeneity: Functional properties of bone marrow-derived and sessile hepatic macrophages. Blood 110, 4077–4085 (2007).

32. Wang, L. xun, Zhang, S. xi, Wu, H. juan, Rong, X. lu & Guo, J. M2b macrophage polarization and its roles in diseases. J. Leukoc. Biol. 106, 345–358 (2019).

33. Ding, B. Sen et al. Divergent angiocrine signals from vascular niche balance liver regeneration and fibrosis. Nature 505, 97–102 (2014).

34. Ding, B. Sen et al. Inductive angiocrine signals from sinusoidal endothelium are required for liver regeneration. Nature 468, 310–315 (2010).

35. Hintermann, E. & Christen, U. The many roles of Cell Adhesion Molecules in Hepatic Fibrosis. Cells 8, (2019).

36. Shubham, S. et al. Cellular and functional loss of liver endothelial cells correlates with poor hepatocyte regeneration in acute-on-chronic liver failure. Hepatol. Int. 13, 777–787 (2019).

37. Yin, K. et al. Senescence-induced endothelial phenotypes underpin immune-mediated senescence surveillance. Genes Dev. 36, 533–549 (2022).

38. Wen, Y., Lambrecht, J., Ju, C. & Tacke, F. Hepatic macrophages in liver homeostasis and diseases-diversity, plasticity and therapeutic opportunities. Cell. Mol. Immunol. 18, 45–56 (2021).

39. Schuppan, D. & Kim, Y. O. Evolving therapies for liver fibrosis. J. Clin. Invest. 123, 1887–1901 (2013).

40. Kocabayoglu, P. et al. β-PDGF receptor expressed by hepatic stellate cells regulates fibrosis in murine liver injury, but not carcinogenesis. J. Hepatol. 63, 141–147 (2015).

41. Wynn, T. A. & Barron, L. Macrophages: Master Regulators of Inflammation and Fibrosis. Semin. Liver Dis. 30, 245–257 (2010).

42. Grier, J. D., Yan, W. & Lozano, G. Conditional allele of mdm2 which encodes a p53 inhibitor. Genesis 32, 145–147 (2002).

43. Wight, T. N. & Potter-Perigo, S. The extracellular matrix: An active or passive player in fibrosis? Am. J. Physiol. - Gastrointest. Liver Physiol. 301, 950–955 (2011).

44. Mcquitty, C. E., Williams, R., Chokshi, S. & Urbani, L. Immunomodulatory Role of the Extracellular Matrix Within the Liver Disease Microenvironment. Front. Immunol. 11, (2020).

