## Supplementary Materials for "Utilising an in silico model to predict outcomes in senescence-driven acute liver injury"

Supplementary Material for: **Utilising an in silico model to predict outcomes in senescence-driven acute liver injury**

### Contents


| | Page number |
|---|---|
| Supplementary Figure 1 | 3 |
| Supplementary Figure 2 | 4 |
| Supplementary Figure 3 | 5 |
| Supplementary Figure 4 | 6 |
| Supplementary Figure 5 | 7 |
| Supplementary Figure 6 | 9 |
| Supplementary Figure 7 | 10 |
| Supplementary Table 1 | 11 |
| Supplementary Table 2 | 12 |
| Supplementary Mathematical Methods | 13 |
| Supplementary Methods | 15 |

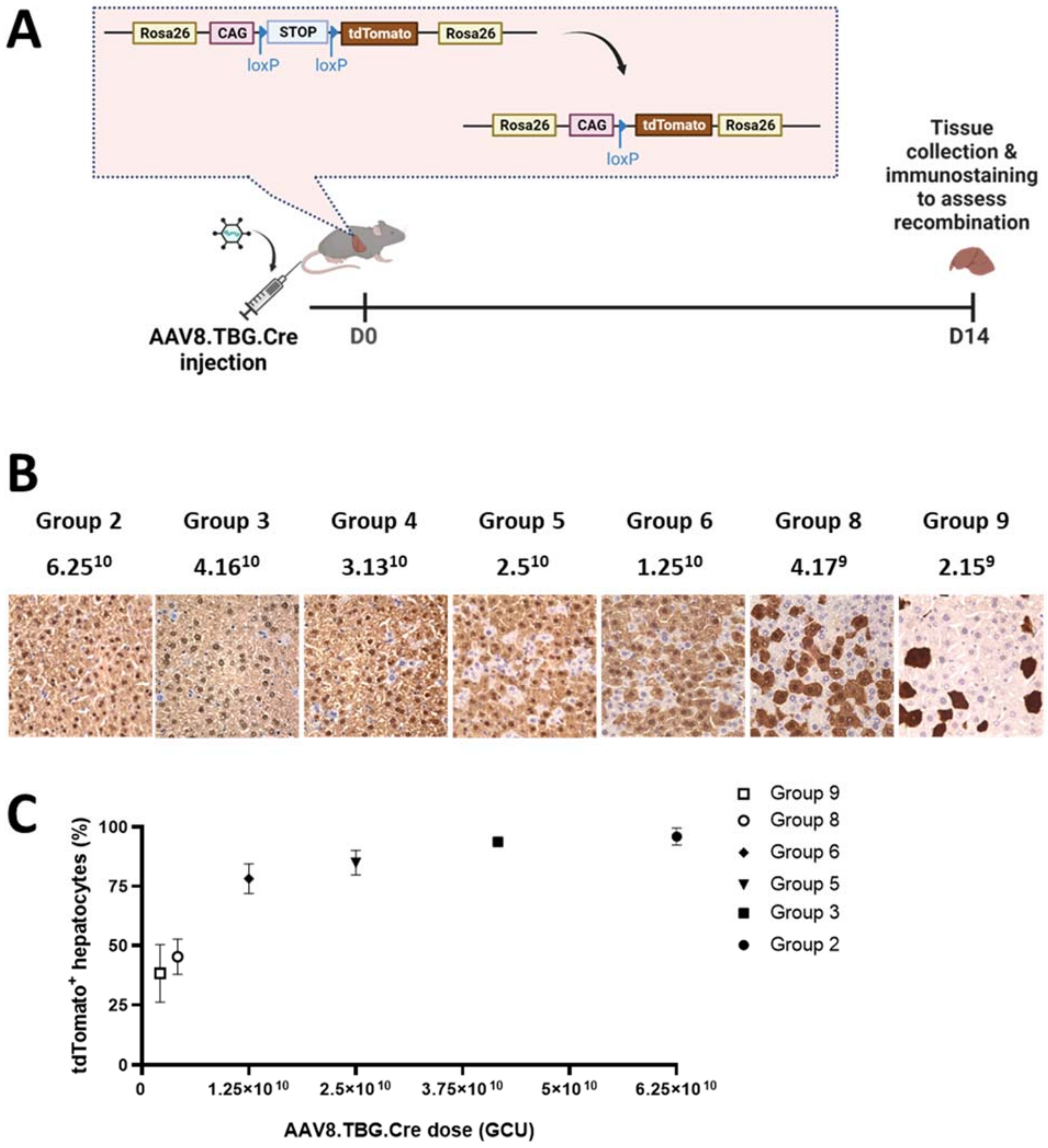


***Supplementary Figure 1- Rosa26$^{LSL\text{-}TdTomato}$ AAV8-TBG-Cre dose response to assess recombination efficiency.*** **[A]-** Graphical schematic for the recombination dose response study. Inset demonstrates tissue specific Cre recombinase expression mediated induction of tdTomato expression in the Rosa26$^{LSL\text{-}TdTomato}$ mouse model. Due to the hepatotropic nature of the AAV8.TBG.Cre, the dose injected impacts the proportion of hepatocytes that express tdTomato as a result of Cre-mediated recombination, allowing assessment of recombination efficiency with different doses based on quantification of the proportion of hepatocytes that are tdTomato positive. **[B]-** Representative image panels from histological sections immunostained for tdTomato demonstrating the changing proportion of positively stained cells. Labels correspond to the equivalent Group in the Mdm2 study (top) and the AAV8 dose in GCU (bottom). **[C]-** Quantification of recombination demonstrating the mean proportion of recombined hepatocytes for each dose as % tdTomato positive cells (N=3 mice per group, N≥5 FOVs analysed per mouse).

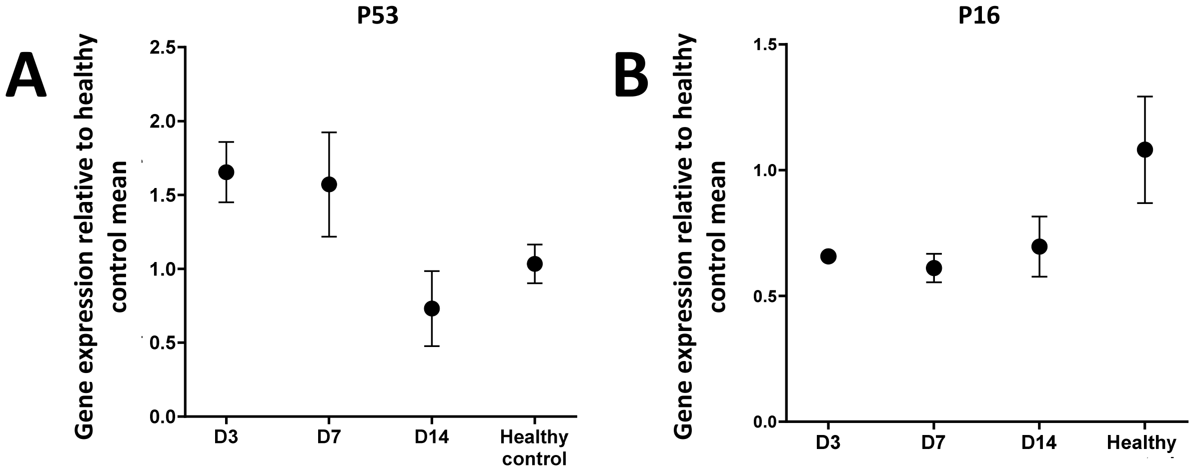


***Supplementary Figure 2- Expression of senescence markers in whole liver tissue of Mdm2 mice administered an AAV8.TBG.Cre dose commensurate to a moderate dose.*** Expression of **[A]-** p53, **[B]-** p16 was analysed at the indicated time points. qPCR results were normalised to PPIA housekeeper and expressed as fold change relative to healthy control mean expression. N=3-5 mice per timepoint analysed. Error bars are SEM. Ordinary One-Way ANOVA relative to healthy control mean with Dunnett's Multiple Comparisons test.

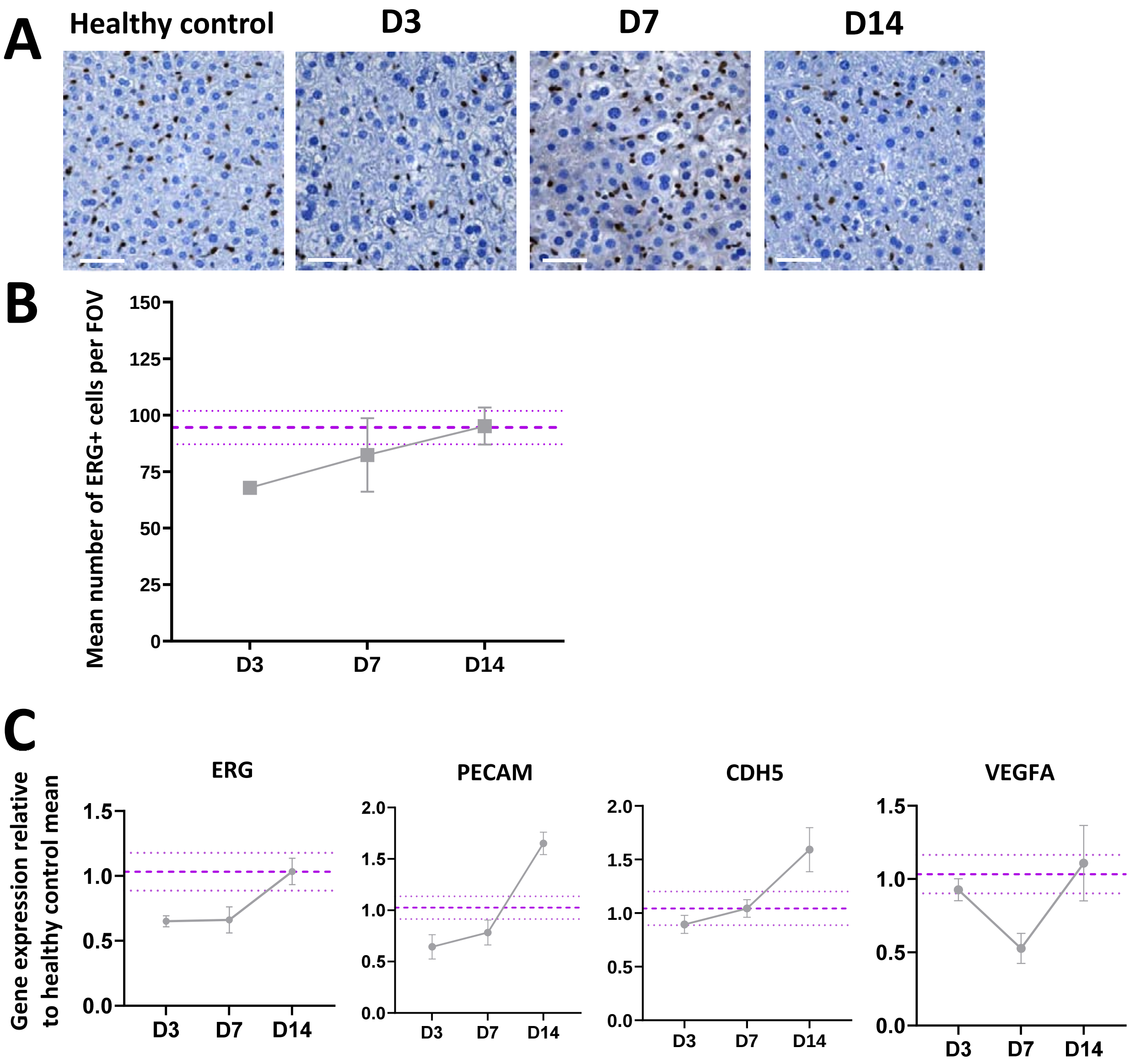


***Supplementary Figure 3- Tissue and gene expression of endothelial markers in whole liver tissue of Mdm2 mice administered an AAV8.TBG.Cre dose commensurate to a moderate dose.***

**[A]-** Representative histological micrographs of ERG (endothelial nuclear cell marker) at D3, D7 and D14 following induction of moderate senescence injury induction. Scale bars are 50µm.

**[B]-** Quantification of immunostaining in [A] as number of positively stained nuclei per FOV. ≥10 FOVs quantified per animal. N=3-5 animals per timepoint

**[C]-** mRNA expression of the endothelial markers ERG, PECAM, CDH5 and VEGFA was analysed at indicated time points. qPCR results were normalised to PPIA housekeeper and expressed as fold change relative to healthy control mean expression. N=3-5 mice per timepoint analysed. Error bars are SEM.

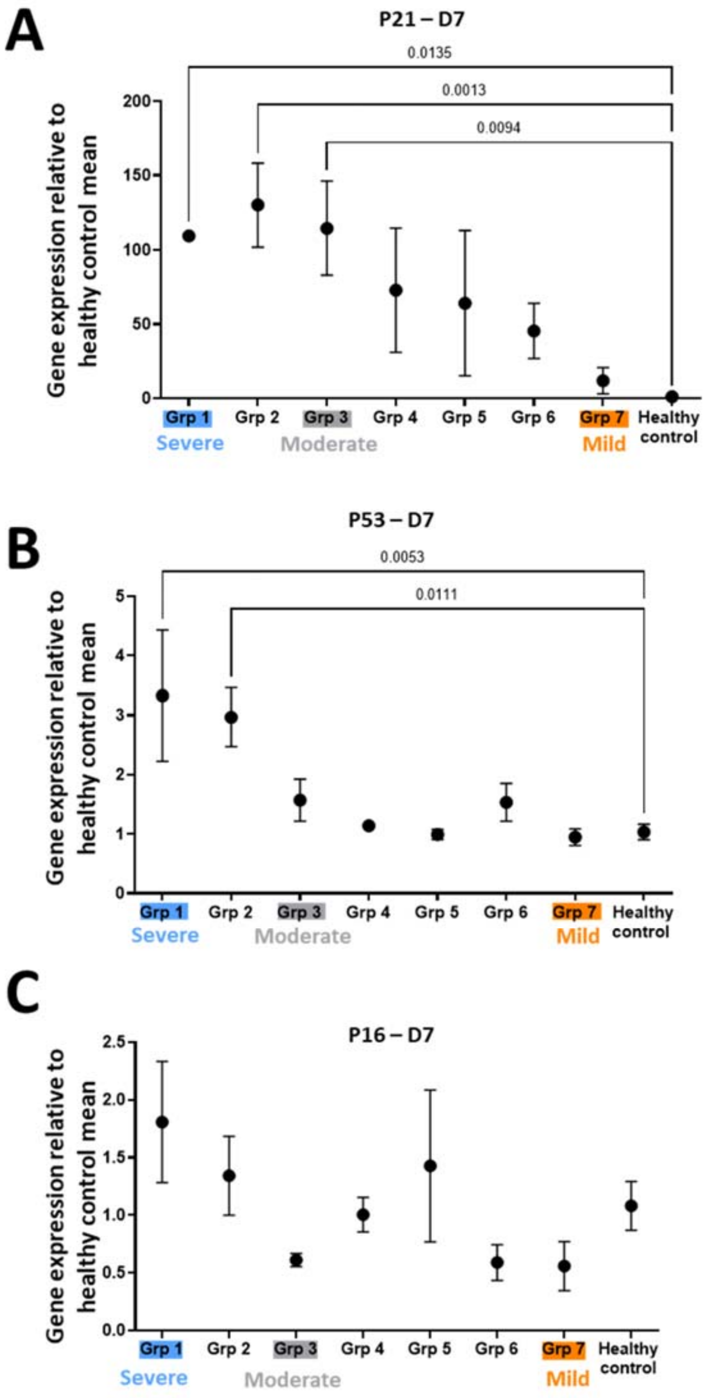


***Supplementary Figure 4- Expression of senescence markers in whole liver tissue of Mdm2 mice administered a range of AAV8.TBG.Cre doses.***

Expression of the senescence markers **[A]** p21, **[B]** p53 and **[C]** p16 across the full senescence dose response analysed at D7 following AAV8.TBG.Cre induction. qPCR results were normalised to PPIA housekeeper and expressed as fold change relative to healthy control mean expression. N=3-5 mice per group analysed. Error bars are SEM. Ordinary One-Way ANOVA relative to healthy control mean with Dunnett's Multiple Comparisons test. Results show dose dependent increases in p21 and p53 expression of these senescent markers but no difference in p16 indicating that AAV8.TBG.Cre induces senescence along the p21/p53 axis.

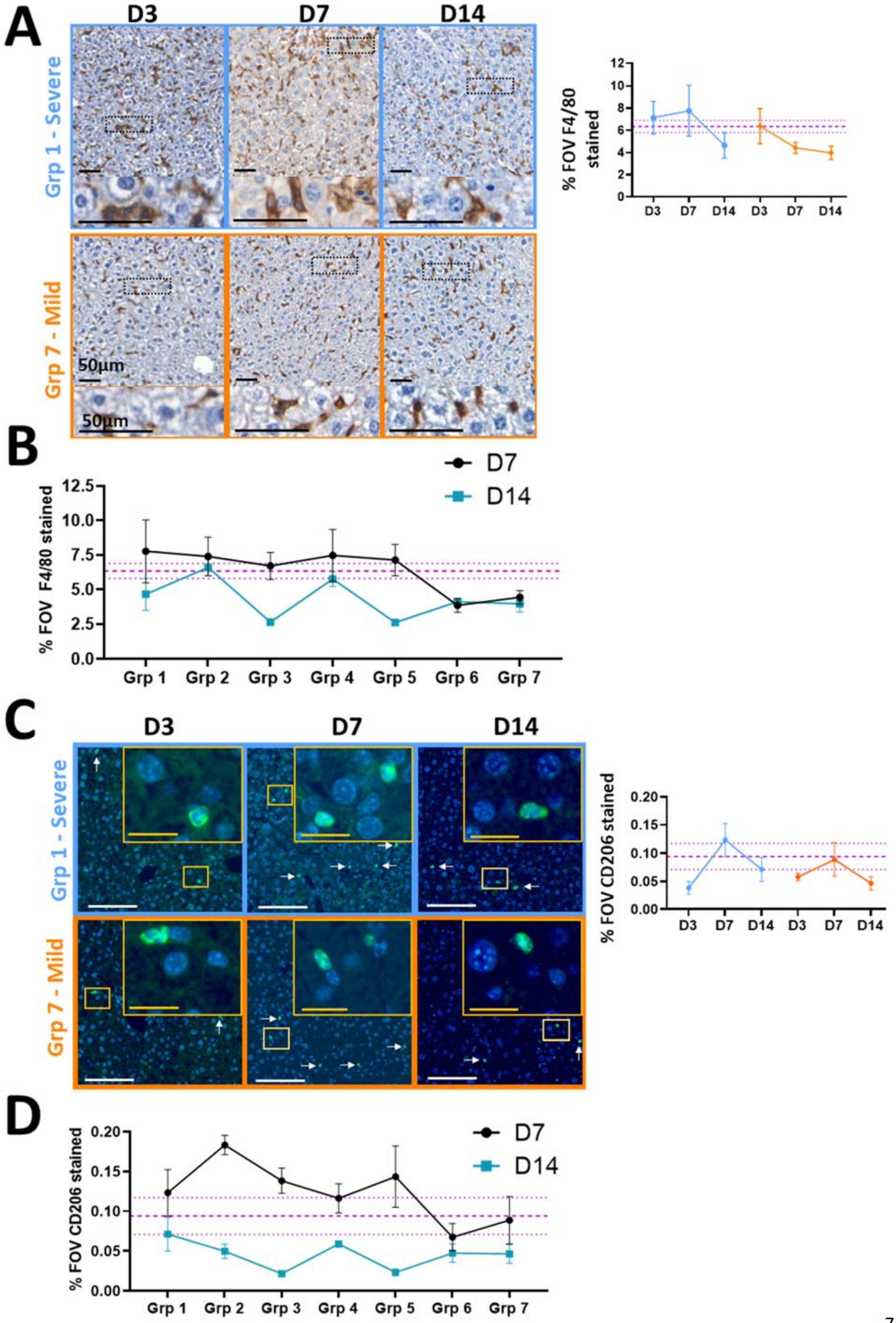
A
D3
D7
D14
Grp 1 - Severe
Grp 7 - Mild
50µm
50µm
% FOV F4/80 stained
B
D7
D14
% FOV F4/80 stained
Grp 1
Grp 2
Grp 3
Grp 4
Grp 5
Grp 6
Grp 7
C
D3
D7
D14
Grp 1 - Severe
Grp 7 - Mild
% FOV CD206 stained
D
D7
D14
% FOV CD206 stained
Grp 1
Grp 2
Grp 3
Grp 4
Grp 5
Grp 6
Grp 7

***Supplementary Figure 5- Quantification results for tissue staining of macrophage markers after induction of different levels of senescence in Mdm2 mice.***

Results of immunohistochemical tissue staining and quantification of Mdm2 mice administered with a range of doses of AAV8.TBG.Cre (group 1-7) to induce different levels of senescence. Quantification of staining based on % of the field of view (FOV) positively stained. N=3-5 independent animals per group and timepoint with ≥10 FOVs quantified per animal. Purple dashed lines are healthy control mean±SEM. **[A]-**Representative histological micrographs of the F4/80 pan macrophage marker in the Severe and Mild senescence injury dosing groups at D3, D7 and D14 post-induction with accompanying quantification. Scale bars 50µm. **[B]-** Quantification of F4/80 staining based on % of FOV at D7 and D14 for the entire dosing cohort. **[C]-** Representative micrographs of CD206 tissue staining in the Severe and Mild senescence injury dosing groups at D3, D7 and D14 post-induction with accompanying quantification. White arrows indicate positively stained cells. Insets are digitally magnified 1.5x. **[D]-** Quantification of CD206 staining based on % of FOV at D7 and D14 for the entire dosing cohort.

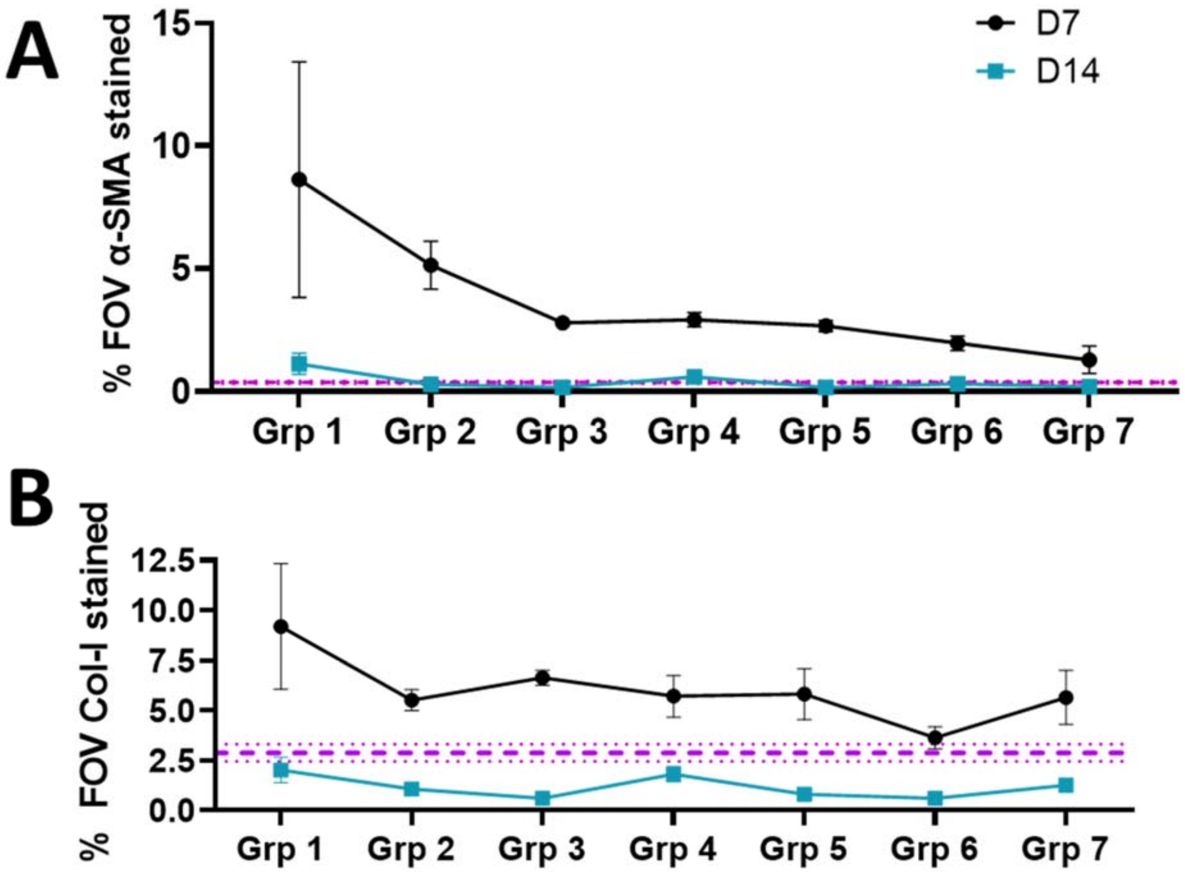


***Supplementary Figure 6- Quantification results for tissue staining of tissue remodelling markers D7 and D14 after induction of different levels of senescence in Mdm2 mice.***

Results of immunohistochemical tissue staining quantification of *Mdm2* mice administered with a range of doses of AAV8.TBG.Cre (group 1-7) to induce different levels of senescence at D7 and D14 **[A]-** α-SMA (activated myofibroblast marker) and **[B]-** Collagen-I (ECM marker). Purple dashed lines are healthy control mean±SEM. Data points represent mean N=3-5 independent animals per group and timepoint. All error bars are ±SEM.

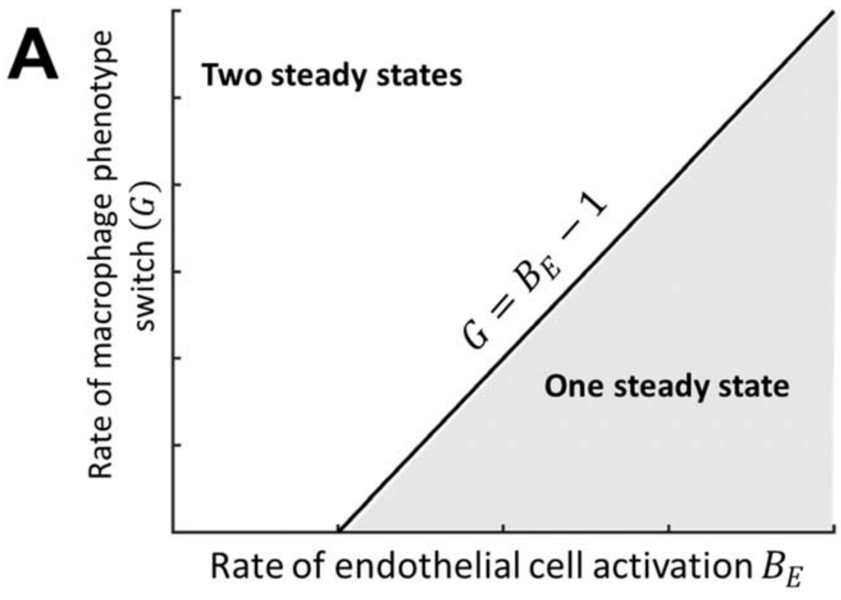


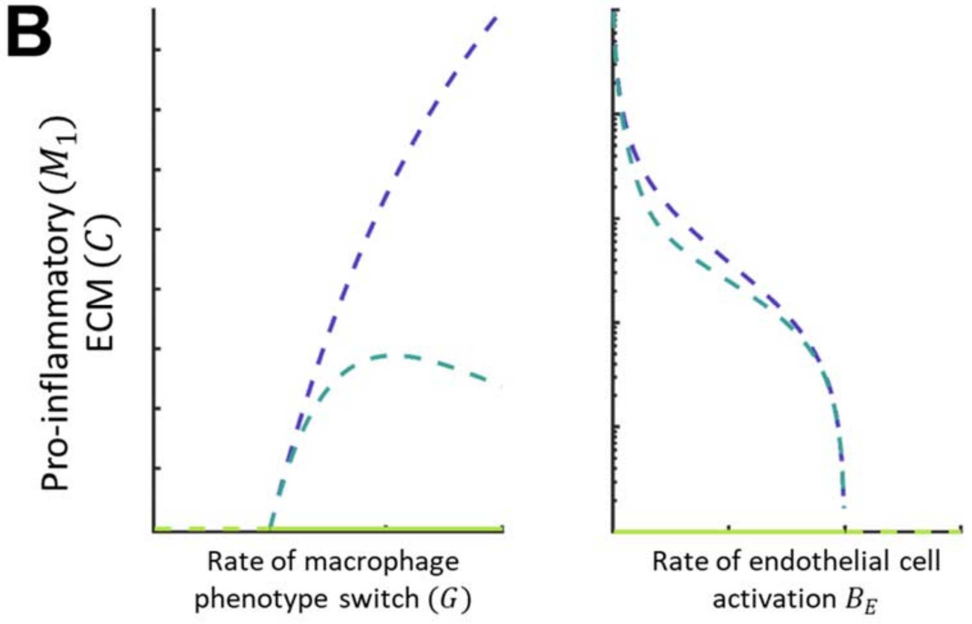


***Supplementary Figure 7- Solution space for rate values and linear stability analysis in bifurcation graph***

**[A]** Solution space and number of steady states for different $G$ and $B_E$.

**[B]** Bifurcation diagrams for pro-inflammatory macrophages $M_1$ (purple line) and ECM $C$ (light blue line) as a function of rate of macrophage phenotype switch $G$ (with $B_E$ set to a value different than 1 and the rest other parameters equal to 1) and rate of endothelial cells activation $B_E$ (all parameters equal to 1). Solid and dashed lines represent stable and unstable solution branches respectively. The green line is the zero steady state. LHS: At a critical value of $G$, a second positive unstable steady branch appears. As $G$ increases, $C$ decreases approaching zero, however it always remains non-zero, positive and unstable. RHS: At a critical value of $B_E$, the two solution branches collapse into one, the zero one, which is unstable.

***Supplementary Table 1- Primary antibody conditions used for immunostaining***

| Antigen | Supplier | Catalogue # | Species | Stock protein conc. (mg/mL) | Dilution | Antigen retrieval |
|---|---|---|---|---|---|---|
| CD206 | Abcam | Ab64693 | Rabbit | 1 | 1/200 | pre-warmed sodium citrate buffer (pH 6.0), 15 mins |
| F4/80 | Abcam | Ab6640 | Rat | 1 | 1/100 | 15 minutes Proteinase K at 37$^{o}$C |
| iNOS | BD | 610329 | Mouse | 0.25 | 1/100 | pre-warmed sodium citrate buffer (pH 6.0), 15 mins |
| α-SMA | Sigma | A2547 | Mouse | 2.0-6.0 (varies) | 1/1000 | 5min pre-heat TE buffer (pH 9.0), 10 mins |
| Col-I | Southern Biotech | 1310-01 | Goat | 0.2 | 1/300 | 5min pre-heat TE buffer (pH 9.0), 10 mins |
| ERG | Abcam | Ab92513<br>Ab214341 | Rabbit<br>Mouse | 0.88 | 1/125<br>1/500 | DAB: pre-warmed sodium citrate buffer (pH 6.0), 10 mins<br>IF: TE 5min pre-warm + 10min |
| VCAM-1 | Abcam | Ab134047 | Rabbit | 0.43 | 1/500 | IF: TE 5min pre-warm + 10min |

***Supplementary Table 2- Details of primers used in qPCR analyses***

| Marker | Gene name | Quantitect # |
|---|---|---|
| Myofibroblast | ACTA2/α-SMA | QT00140119 |
| Myofibroblast | PDGFRB | QT00113148 |
| Inflammation | SOCS3 | QT02488983 |
| Tissue macrophage/Kupffer cells | CD68 | QT00254051 |
| Tissue macrophage/Kupffer cells | EMR1/F4/80 | QT00099617 |
| M2 macrophage | MRC1/CD206 | QT00103012 |
| M2 macrophage | CD163 | QT00123074 |
| M1 macrophage | CD86 | QT01055250 |
| M1 macrophage | CD80 | QT00129787 |
| Collagen ECM marker | Collagen 1A1 | QT02589482 |
| Collagen ECM marker | Collagen 1A2 | QT02325736 |
| Endothelial/endothelial proliferation | PECAM-1/CD31 | QT01052044 |
| Endothelial/endothelial proliferation | VEGFA | QT00160769 |
| Endothelial/endothelial proliferation | ERG | QT01755551 |
| Endothelial/endothelial proliferation | CDH5/VE-Cadherin | QT00110467 |
| Endothelial/endothelial proliferation | FGA | QT00128198 |
| Endothelial/endothelial proliferation | FGG | QT00101738 |
| Endothelial/endothelial proliferation | VCAM-1 | QT00128793 |
| Acute endothelial injury/regeneration marker | ACKR3/CXCR7 | QT00254443 |
| Endothelial activation marker | ICAM-1 | QT00155078 |
| Senescence | CDKN2A/P16 | QT00252595 |
| Senescence | CDKN1A/P21 | QT00137053 |
| Senescence | TRP53/P53 | QT00101906 |
| Housekeeper | PPIA | QT00247709 |

**Supplementary Mathematical Methods**

**Steady states**

Steady state solutions (that have no time dependence) can be determined by setting the left hand side of the equations corresponding to time derivatives to zero (1a-1f). We denote steady states variables with stars. We note that equations (1a)-(1f) have a zero steady state $(T^*, M_1^*, M_2^*, E^*, C^*, F^*) = (0,0,0,0,0,0)$. To determine the non-zero steady states of the model, we set the right hand sides of (1a)-(1f) to zero and solve the resulting coupled algebraic equations. We reduce the system to two non-linear algebraic equations for F* and C* as follows:

$$F^* = \frac{B_F f(y^*)}{D + K_F g(y^*)}, C^* = \frac{B_C B_F f(y^*)}{[D + K_C g(y^*)]\,[D + K_F g(y^*)]} \qquad \text{Equation (3)}$$

where

$$f(y^*) = \frac{D}{B_E} - \frac{y^*}{Dy^* + G}, g(y^*) = \frac{G}{D}\frac{f(y^*)}{y^*}, y^* = (1 + C^*)(1 + F^*) \qquad \text{Equation (4)}$$

The steady state values of the remaining variables are then given by

$$T^* = 0, M_1^* = f(y^*), M_2^* = g(y^*), E^* = \frac{B_E f(y^*)}{D - B_E f(y^*)} \qquad \text{Equation (5)}$$

We can use these results to determine conditions on the parameters to obtain the non-zero steady state. Since $C^*$and $F^*$ are positive, we have that $y^* > 1$ and therefore

$f >= \frac{D}{B_E} - \frac{1}{D+G}$ and Equation (6)

$$g > \frac{G(D^2 + DG - B_E)}{B_E(D^2 + DG)} \qquad \text{Equation (7)}$$

Since we also require $f > 0$ and $g > 0$ to ensure positive macrophage populations, we therefore find

$$G > \frac{B_E - D^2}{D} \qquad \text{Equation (8)}$$

which is the condition on the parameters for two steady state solutions to co-exist.

**Linear stability analysis**

To determine the linear stability of the steady state solutions we introduce small amplitude perturbations of the form

$$T = \mathrm{T} + \varepsilon \hat{T} E^{\lambda t} + \mathrm{O}(\varepsilon^2) \qquad \text{Equation (9)}$$

With similar expressions for $M_1, M_2, E, C$ and $F$. Here λ is the growth rate of the perturbation, $\hat{T}$ etc are constants and $0 < \varepsilon \ll 1$ is a small parameter. Substituting the expressions (9)_ into equations (1a)-(1f) and retaining linear terms leads to an equation of the form

$$\boldsymbol{J}x = \lambda x \qquad \text{Equation (10)}$$

where $\boldsymbol{J}(T^*, M_1^*, M_2^*, E^*, C^*, F^*)$ is the 6X6 Jacobian matrix given by

$$\begin{bmatrix} -K_T M_2^* - D & 0 & -K_T T^* & 0 & 0 & 0 \\ B_1 & -\dfrac{G}{(C^*+1)(F^*+1)} - D & 0 & 1 & \dfrac{GM_1^*}{(C^*+1)^2(F^*+1)} & \dfrac{GM_1^*}{(C^*+1)(F^*+1)^2} \\ B_2 & \dfrac{G}{(C^*+1)(F^*+1)} & -D & 0 & -\dfrac{GM_1^*}{(C^*+1)^2(F^*+1)} & -\dfrac{GM_1^*}{(C^*+1)(F^*+1)^2} \\ 0 & B_E(1+E^*) & 0 & B_E M_1^* - D & 0 & 0 \\ 0 & 0 & -K_C C^* & 0 & -K_C M_2^* - D & B_C \\ 0 & B_F & -K_F F^* & 0 & 0 & -K_F M_2^* - D \end{bmatrix}$$

Equation (10), $\boldsymbol{x}$ is the vector of unknowns $\boldsymbol{x} = (\hat{T}, \widehat{M}_1, \widehat{M}_2, \hat{E}, \hat{C}, \hat{F})$ and the eigenvalues $\lambda = \lambda_n$ $(n = 1, \dots, 6)$ satisfy $|\boldsymbol{J} - \lambda \boldsymbol{I}| = 0$. Instability of a steady state is predicted when $\max\limits_{\{n=1,\dots,6\}} R(\lambda_n) > 0$. Conversely, the steady state is said to be linear stable if $R(\lambda_n) < 0$ for $n = 1, \dots, 6$ where $R$ is the real part of the eigenvalues.

The eigenvalues associated with the trivial steady state are real numbers. The largest eigenvalue is given by

$$\lambda = \frac{1}{2}\left(-2D - G + \sqrt{4B_E + G^2}\right), \qquad \text{Equation (11)}$$

and we see that the stability of the trivial steady state depends only on $G$, the rate of phenotype switch and $B_E$, the rate of activated endothelial cell increase, and $D$. This eigenvalue is negative when

$$B_E > 0, G > max\left\{0, \frac{B_E - D^2}{D}\right\} \qquad \text{Equation (12)}$$

which ensures the origin is stable point. This condition is also required for the existence of the non-zero steady state.

A similar linear stability of the non-trivial steady state indicates that it is always unstable to small amplitude perturbations (details not shown).

In **Supplementary Figure 7**, we present bifurcation diagrams showing how the steady-state solution structure varies as $B_E$ and $G$ are varied. All other parameters are set to 1. The transcritical bifurcation point corresponds to $G = B_E - 1$. In the left hand figure, as the parameter $G$ decreases through the bifurcation point, the non-trivial steady state is lost and there is a change in stability for the trivial steady state from stable to unstable. A similar behaviour is observed in the right hand figure, as $B_E$ increases through the bifurcation point.

#### Supplementary Methods

***Supplementary Method 1- Representative Fiji macro for automated quantification of stained regions***

```
/* Macro for quantifying area of DAB staining in haemotoxylin and DAB stained tissues
 */

//CLEAR LOG
print("\\Clear");

// CLOSE ALL OPEN IMAGES
while (nImages>0) {
		selectImage(nImages);
   close();
}

//SET BATCH MODE
setBatchMode(false);   //true or false, true if you don't want to see the images, which is faster

//START MESSAGE
print("**** STARTING THE MACRO ****");

//INPUT/OUPUT folders
inDir=getDirectory("Choose the input folder");
outputDir=getDirectory("And the output folder");
myList=getFileList(inDir);  //an array

//Define your measurements and settings for
run("Set Measurements...", "area area_fraction limit redirect=None decimal=2");
roiManager("Set Line Width", 2);

//Defining arrays
ImageName=newArray();
PercentageSurfaceInDAB=newArray();
AreaInDAB=newArray();
ImageAnalysed=0;

for (j = 0 ; j < myList.length ; j++ ){
		path=inDir+myList[j];   //path to each file
		open(path);
		FileName=File.nameWithoutExtension;
		ImageID=File.name;
		Title=getTitle();
		print("Processing "+ImageID);
		ImageName=Array.concat(ImageName,Title);
		getStatistics(area, mean, min, max, std, histogram);


		//Thresholding the image to find DAB.  First split image by Colour Deconvolution and keep DAB
channel
		selectWindow(Title);
		run("Set Scale...", "distance=3.9924 known=1 unit=µm global");
```

```
            run("Duplicate...", "title=DAB");
            run("Colour Deconvolution", "vectors=[H DAB]");
            selectWindow("DAB-(Colour_3)");
            close();
            selectWindow("DAB-(Colour_1)");
            close();
            selectWindow("Colour Deconvolution");
            close();
            selectWindow("DAB-(Colour_2)");

            //set threshold for DAB positive regions (modify depending on stain isotype control)
            run("Threshold...");
            setThreshold(0, 180);
            setOption("BlackBackground", true);
            run("Convert to Mask");

            //convert to binary image
            run("Measure");


            //updating area results
            AreaDAB=getResult("Area",j);
            AreaInDAB=Array.concat(AreaInDAB,AreaDAB);
            PercentAreaDAB=(AreaDAB/area)*100;
            PercentageSurfaceInDAB=Array.concat(PercentageSurfaceInDAB,PercentAreaDAB);


            //QC images with mask overlay
            selectWindow("DAB-(Colour_2)");
            run("Analyze Particles...", "size=0-Infinity add");
            selectWindow("DAB");
            roiManager("Show All without labels");
            RoiManager.setPosition(0);
            roiManager("Set Color", "yellow");
            roiManager("Set Line Width", 2);
            run("Flatten");

//saving QC image
            selectWindow("DAB-1");
            saveAs("Tiff", outputDir+Title+"_QC.tif");
            close();


            print("There was an area of "+AreaDAB+" stained, representing "+PercentAreaDAB+" percent of the
FOV");

            close("*");

            roiManager("reset");
}

Array.show("Results", ImageName,AreaInDAB,PercentageSurfaceInDAB);
            saveAs("Results", outputDir+"Results.csv");
```

```
//saving log
print("***** Macro done *****");
selectWindow("Log");
saveAs("Text", outputDir+FileName+"_Log.txt");

close("*");
```

***Supplementary Method 2- Representative Fiji macro for automated quantification of stained nuclei (ERG analysis)***

```
/* Macro for quantifying area of DAB staining in haemotoxylin and DAB stained tissues
 */

//CLEAR LOG
print("\\Clear");

// CLOSE ALL OPEN IMAGES
while (nImages>0) {
		selectImage(nImages);
   close();
}

//SET BATCH MODE
setBatchMode(false);   //true or false, true if you don't want to see the images, which is faster

//START MESSAGE
print("**** STARTING THE MACRO ****");

//INPUT/OUPUT folders
inDir=getDirectory("Choose the input folder");
outputDir=getDirectory("And the output folder");
myList=getFileList(inDir);  //an array

//Define your measurements and settings for
run("Set Measurements...", "area area_fraction limit redirect=None decimal=2");
roiManager("Set Line Width", 2);

//Defining arrays
ImageName=newArray();
PercentageSurfaceInDAB=newArray();
AreaInDAB=newArray();
DABPatches=newArray();
ImageAnalysed=0;

for (j = 0 ; j < myList.length ; j++ ){
		path=inDir+myList[j];   //path to each file
		open(path);
		FileName=File.nameWithoutExtension;
		ImageID=File.name;
```

```
	Title=getTitle();
	print("Processing "+ImageID);
	ImageName=Array.concat(ImageName,Title);
	getStatistics(area, mean, min, max, std, histogram);


	//Thresholding the image to find DAB.  First split image by Colour Deconvolution and keep DAB
channel
	selectWindow(Title);
	run("RGB Color");
	run("Duplicate...", "title=DAB");
	run("Split Channels");
	selectWindow("DAB (green)");
	selectWindow("DAB (blue)");
	selectWindow("DAB (red)");
	close();
	selectWindow("DAB (green)");
	close();
	setAutoThreshold("Default dark");

	//run("Threshold...");
	setThreshold(150, 255);
	run("Invert");
	run("Analyze Particles...", "size=300-Infinity pixel add");
	DAB=roiManager("count");


	//Updating array to include the number of patches
	DABPatches=Array.concat(DABPatches,DAB);


	//Circumventing issues if there is no or only one DAB patch by creating single pixel ROIs
	if(DAB<=2){
		makeRectangle(0, 0, 1, 1);
		roiManager("Add");
		makeRectangle(1, 1, 1, 1);
		roiManager("Add");
		DAB=DAB+2;
	}

	selectWindow(Title+" (RGB)");
	roiManager("Show All without labels");
	roiManager("Combine");
	roiManager("Add");
	roiManager("Set Color", "yellow");
	roiManager("Set Line Width", 2);
	roiManager("Select All");
	roiManager("Measure");
	run("Flatten");



	//updating area results
```

```
            AreaDAB=getResult("Area",j);
            AreaInDAB=Array.concat(AreaInDAB,AreaDAB);
            PercentAreaDAB=(AreaDAB/area)*100;
            PercentageSurfaceInDAB=Array.concat(PercentageSurfaceInDAB,PercentAreaDAB);


            //QC images with mask overlay

            selectWindow(Title+" (RGB)-1");
            saveAs("Tiff", outputDir+Title+"_QC.tif");
            close();

            print("There were "+DAB+" ERG stained cells, with an area of "+AreaDAB+" stained, representing
"+PercentAreaDAB+" percent of the FOV");

            close("*");

            roiManager("reset");
}

Array.show("Results", ImageName,DABPatches,AreaInDAB,PercentageSurfaceInDAB);
            saveAs("Results", outputDir+"Results.csv");


//saving log
print("***** Macro done *****");
selectWindow("Log");
saveAs("Text", outputDir+FileName+"_Log.txt");

close("*");
```
